# Mast cell-driven adipocyte lipolysis promotes post-expansive VAT atrophy and ectopic hepatic steatosis

**DOI:** 10.64898/2026.01.07.698076

**Authors:** Jiahui Wang, Lei Zhang, Wen-Yue Liu, Xin-Zhe Jin, Hao Yin, Hao Cai, Hai-Yang Yuan, Wah Yang, Bin Bao, Ming-Hua Zheng, Hua Wang, Guo-Ping Shi, Xian Zhang, Jian Liu

## Abstract

Excessive visceral fat is an essential risk factor of metabolic dysfunction-associated steatotic liver disease (MASLD). Yet, the cellular and molecular mechanisms implicated in the correlation between visceral fat metabolism and hepatic steatosis remain poorly understood. Here we report post-expansive epididymal adipose tissue (EAT) atrophy and hepatic steatosis in mice fed a high-fat diet (HFD) primarily due to elevated adipocyte lipolysis in EAT. Mast cell (MC) accumulation in EAT represents a lipolysis-associated feature. Pharmacological stabilization of MCs suppresses EAT adipocyte lipolysis, and improves EAT atrophy and hepatic steatosis. MC-derived serotonin (5-HT) correlates with visceral adipose tissue (VAT) lipolysis in HFD-fed mice as well as in MASLD patients. Conditional deletion of 5-HT from MCs or its receptor HTR2b on adipocytes demonstrates that MC-derived 5-HT promotes adipocyte lipolysis, EAT atrophy, and hepatic steatosis by binding on adipocyte HTR2b. These results suggest that MCs and MC-derived 5-HT are potential therapeutic targets for obesity-associated MASLD.

## Introduction

Obesity and associated adipose tissue remodeling is correlated with the development of several metabolic complications, including nonalcoholic fatty liver diseases (NAFLD)^1-3^, a metabolic dysfunction-associated fatty liver disease or metabolic dysfunction-associated steatotic liver disease (MAFLD/MASLD)^4,5^. Yet, not all obese individuals suffer from MAFLD/MASLD and not all fat accumulation in adipose tissue contributes to these common metabolic diseases. Clinical, epidemiologic, and experimental investigations suggest that abdominal obesity and excessive visceral fat, but not subcutaneous obesity and excessive subcutaneous fat are risk factors of MAFLD/MASLD^6-9^. In mice fed a high-fat diet (HFD), body weight gain and excessive energy intake are accompanied with sustained expansion of subcutaneous adipose tissue (SAT)^8^. Unlike SAT, the expansion of epididymal adipose tissue (EAT), which refers to visceral adipose tissue (VAT) occurs in a distinct process. EAT expands first and then undergoes atrophy^8^. Meanwhile, loss of EAT depot is inversely correlated with hepatic steatosis, suggesting an ectopic fat relocation into liver from EAT^8^. However, the detailed cellular and molecular interplays between post-expansive EAT lipoatrophy and ectopic hepatic steatosis remain unknown.

White adipose tissue (WAT) is an endocrine organ essential for energy metabolism and immunoregulation^10,11^. In WAT, white adipocytes are differentiated from pre-adipocytes or adipocyte progenitors in the stromal vascular fraction (SVF). Mature adipocytes not only store energy *via* lipogenesis in the form of triglycerides (TG), but also convert TG into non-esterified fatty acids (NEFA, also known as free fatty acids FFA) and glycerol through lipolysis for energy mobilization. Thus, adipocyte size and turnover determine WAT mass^10,12^. In mice fed a HFD, adipose tissues expand *via* adipocyte hypertrophy (increase in size) and hyperplasia (increase in number) in response to overnutrition^10,12,13^. Besides pre-adipocytes and adipocyte progenitors, SVF also contains immune cells and endothelial cells (ECs). In mice on a chow diet (CD), EAT maintains in a relatively anti-inflammatory immune state and composes of diverse populations of resident immune cells. After diet-induced obesity (DIO), hypertrophic adipocytes secrete chemokines to recruit pro-inflammatory cells, including mast cells (MCs) into EAT^10,14,15^. Among these pro-inflammatory cells, macrophages are the most abundant and best studies. Macrophage accumulation leads to excessive secretion of pro-inflammatory cytokines and the formation of crown-like structures (CLS), ultimately resulting in adipocyte insulin resistance, dysfunction, and death^16,17^. Healthy adipose tissue expansion involves coordinated EC proliferation and vasculature network growth for nutrients, oxygen, and hormones supplies that are essential for adipocyte survival and function^18-20^. During adipocyte hypertrophy and pro-inflammatory cell infiltration however, EC proliferation and neovasculature growth do not coordinate with adipose tissue expansion, resulting in local persistent hypoxia and even adipocyte death^21,22^. To date, these prior studies mostly remain observational.

A growing body of evidences suggest a role for MCs in various physiological and pathological conditions^23,24^. The numbers of tryptase^+^ or tryptase^+^ chymase^+^ MCs in WAT are higher in obese patients than in lean individuals^25,26^. When mice consumed a high-cholesterol western diet (WD), they gained more body weight with elevated insulin resistance and EAT MC accumulation than those on a CD. MC stabilization with disodium cromoglycate (DSCG) efficiently blocked these pathological changes^25,27^. In mice fed a HFD that contains no cholesterol however, DSCG did not affect body weight gain or insulin sensitivity, although MC accumulation still occurred in EAT^27-30^. Mechanistic studies revealed that dietary cholesterol elevated plasma low-density lipoprotein-cholesterol (LDL) level, resulting in EAT MC activation^27^.

In this study, we fed mice a cholesterol-free HFD and detected a time-dependent EAT expansion (0 to 10 weeks) and atrophy (12 to 20 weeks) and post-expansive hepatic steatosis. Using this model, we tested a role for MCs in post-expansive EAT atrophy and hepatic steatosis. MC-derived 5-HT correlated with VAT lipolysis in HFD-fed mice and MASLD patients. MCs release serotonin (5-HT) to bind its receptor HTR2b on adipocyte and promote adipocyte lipolysis *via* the SIRT1/AMPKα pathway. We used both MC-selective 5-HT-deficient mice and adipocyte-selective *Htr2b*-deficient mice to confirm a critical role for MC-adipocyte interplays in regulating the pathophysiological events implicated in obesity-associated MASLD.

## Results

### 1. Elevated adipocyte lipolysis contributes to EAT atrophy and associated hepatic steatosis in HFD-fed mice

To investigate the dynamic cellular and molecular changes during EAT remodeling, we fed 5-week-old male C57BL/6N wild-type (WT) mice a CD or HFD. HFD-fed mice gained much higher body weight and metabolic tissue mass than CD-fed control mice (Extended Data Fig. 1a-1e). Different from the continuous increases in SAT and brown adipose tissue (BAT) mass, EAT mass increased gradually during the first 10 weeks but reduced from week 12 to 20 (Fig. 1a and Extended Data Fig. 1d/1f). Meanwhile, liver weight remained unchanged from week 0 to 8 and but progressively enhanced from week 10 to 20 (Fig. 1b and Extended Data Fig. 1e/1f), and liver weight gain correlated inversely with EAT weight loss from week 12 to 20 (Fig. 1c). Along with the progressive increase in liver weight and liver fat cavitation area (Fig. 1d and Extended Data Fig. 1g), the expression of genes involved in hepatic lipid transport and synthesis (such as *Scd1*, *CD36*, *Fas*, and *Me1*) was upregulated, whereas the master transcriptional regulator *Pparα* that controls hepatic fatty acid β-oxidation and ketogenesis was downregulated in livers (Extended Data Fig. 1h/1i). Therefore, there were two phases of EAT mass changes in HFD-fed mice, including EAT expansion from week 0 to 10 and EAT atrophy from week 12 to 20. Post-expansive EAT atrophy correlated with ectopic hepatic steatosis.

**Figure 1.**
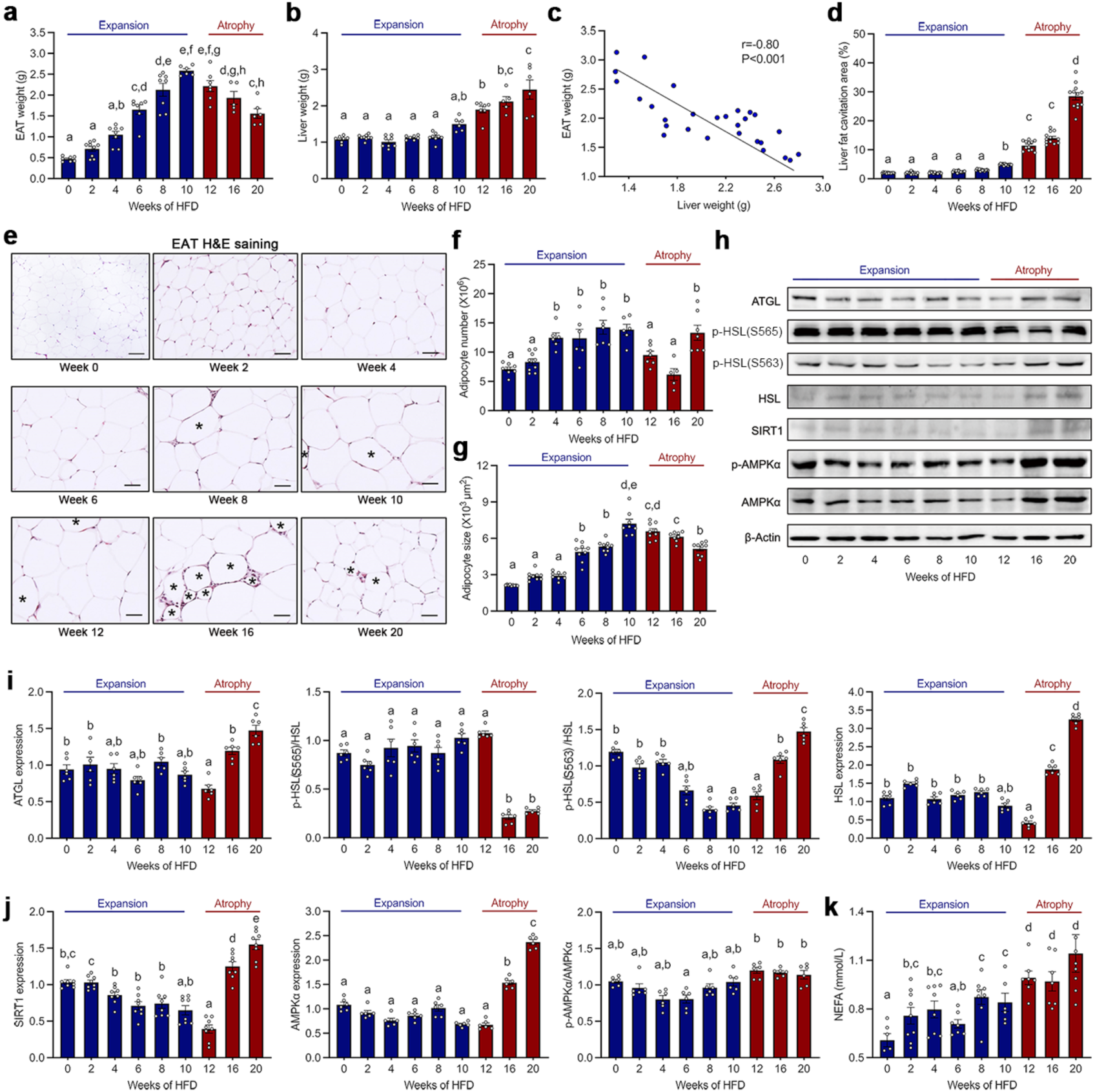
Elevated adipocyte lipolysis contributes to EAT atrophy and associated hepatic steatosis in HFD-fed mice. (**a/b**) EAT weight (**a**) and liver weight (**b**) in WT mice fed an HFD for 20 weeks. (**c**) Correlations between EAT weight and liver weight during EAT atrophy in HFD-fed mice. Pearson’s correlation test. P < 0.05 was considered significant correlation. (**d-g**) Percentage of liver fat cavitation area (**d**); representative images of H&E staining (**e**), adipocyte number (**f**), and adipocyte size (**g**) in EAT in HFD-fed WT mice. Scale bars: 50 µm. Asterisks (*) indicate the CLS. (**h-j**) Immunoblot analysis (**h**) and quantification of (**i**) ATGL and HSL relative to β-Actin, p-HSL(565) and p-HSL(563) relative to HSL, (**j**) SIRT1 and AMPKα relative to β-Actin, and p-AMPKα relative to AMPKα in EAT in HFD-fed WT mice. (**k**) Serum NEFA levels in HFD-fed WT mice. Data are mean ± SEM, n=6-8/ea. Kruskal-Wallis H test with Dunn’s post hoc adjustment for multiple testing, *p* < 0.05 was considered significant. Different letters indicate statistically significant difference.

Adipocyte size and number are the major determinants of WAT mass^10,12^. Progressive upregulation of adipogenic transcription factor *Srebp1*, and the gradual downregulation of cell proliferation gene *E2f1*, adipogenic transcription factors *PPARγ* and *C/EBPα* (Extended Data Fig. 2a), and mature adipocyte markers *Perilipin*, *Adiponectin*, and *Leptin* (Extended Data Fig. 2b) during EAT atrophy in HFD-fed mice. The initial increase and subsequent decrease in cell death during EAT remodeling were demonstrated by TUNEL staining and immunoblot analyses of apoptosis and necrosis markers caspase 3 and RIP3 (Extended Data Fig. 2c-2f). These observations suggest that EAT atrophy is linked to adipocyte differentiation, dysfunction, and death.

Same as reported previously^8^, we found that EAT adipocyte number gradually increased during EAT expansion and rapidly declined at weeks 12 and 16, but then the number was restored at week 20 to the level of week 10 (Fig. 1e/1f). Consistent with the dynamical change of EAT mass, epididymal adipocyte size also gradually enlarged during EAT expansion, following a sudden reduction of adipocyte size during EAT atrophy (Fig. 1e/1g and Extended Data Fig. 2g). The sustained reduction in adipocyte size during EAT atrophy and the comparable level of adipocyte number between week 10 and 20 suggests that the EAT atrophy is primarily determined by the decline in adipocyte size, rather than reduced adipocyte number. In other words, an altered balance between adipocyte lipogenesis and lipolysis, but not between adipocyte differentiation and death, is a major contributor to lipid and weight loss during EAT atrophy.

We assessed the mRNA levels of genes involved in lipogenesis and lipolysis. In HFD-fed mice, although *Dgat1* level was continuously downregulated, all other triglyceride synthetic enzymes *Dgat2*, *Gpat3*, and *Mogat2* were not affected throughout the EAT remodeling (Extended Data Fig. 3a). Unlike the stable lipogenesis maintained by *Dgat2*, *Gpat3*, and *Mogat2*, the expression of lipolytic enzymes *Atgl*, *Hsl*, and *Mgl* (Extended Data Fig. 3b) was enhanced during EAT atrophy, suggesting that the EAT mass and lipid loss depend on the elevated lipolysis but not the inhibited lipogenesis. To test this notion, we detected the protein levels of ATGL, phosphorylated-HSL at S563 (p-HSL(S563)) with lipolytic activity or at S565 (p-HSL(S565)) with anti-lipolytic activity, and total HSL. Relative to the overall stable fluctuation during EAT expansion, the protein levels of ATGL, HSL, and p-HSL(S563) were progressively elevated during EAT atrophy and the p-HSL(S565) level was sharply reduced at weeks 16 and 20 of HFD (Fig. 1h/1i). *Atgl* transcription is upregulated through AMPK and SIRT1 pathways, and inhibited by G0S2^31^. Consistent with increased lipolysis during EAT atrophy, mRNA and protein expression of *Sirt1* and *Ampkα* were also progressively increased, but the mRNA expression of *G0s2* was gradually declined (Fig. 1h/1j and Extended Data Fig. 3c). Abnormal lipoprotein metabolism is closely linked with obesity-associated hepatic steatosis and increased NEFA level from VAT lipolysis is causally involved in the pathophysiology of ectopic fat deposition in patients with visceral obesity^1,32^. Different from the overall stable fluctuation levels of TG and high-density lipoprotein-cholesterol, HFD led to persistent elevation of serum total cholesterol, LDL and NEFA levels (Fig. 1k and Extended Data Fig. 3d-3g). Serum NEFA levels during EAT atrophy were higher than those during EAT expansion (Fig. 1k).

Together, reduced EAT adipocyte size, enhanced EAT lipolysis, and elevated serum NEFA level during EAT atrophy suggest that adipocyte lipolysis serves as the primary cause of EAT lipid loss to promote EAT atrophy and associated hepatic steatosis in HFD-fed mice.

### 2. MC accumulation represents a lipolysis-associated feature during EAT atrophy

Angiogenesis is a rate-limiting step for adipose tissue expansion and remodeling, and plays a key causal role in adipose tissue inflammation, adipocyte dysfunction and death^18,20,22^. In HFD-fed mice, supporting the marked EAT reddening at week 20 relative to week 0 and week 10 (Extended Data Fig. 1f), CD31 immunostaining showed that the angiogenic level remained stable during EAT expansion but dramatically increased during EAT atrophy (Fig. 2a and Extended Data Fig. 4a). Similar to the dynamical change in CLS number in EAT (Fig. 1e and Extended Data Fig. 4b), immunostaining of Mac-2 and real-time PCR analysis of total macrophage marker *F4/80* and M1-like macrophage marker *CD11c* showed that HFD led to sustained macrophage accumulation with peak infiltration observed between week 12 to week 16 (Extended Data Fig. 4c-4f). Therefore, EAT angiogenic activity was characteristically increased during EAT atrophy, and the dynamic infiltrations of both total and pro-inflammatory macrophages were not synchronized with increased adipocyte lipolysis.

**Figure 2.**
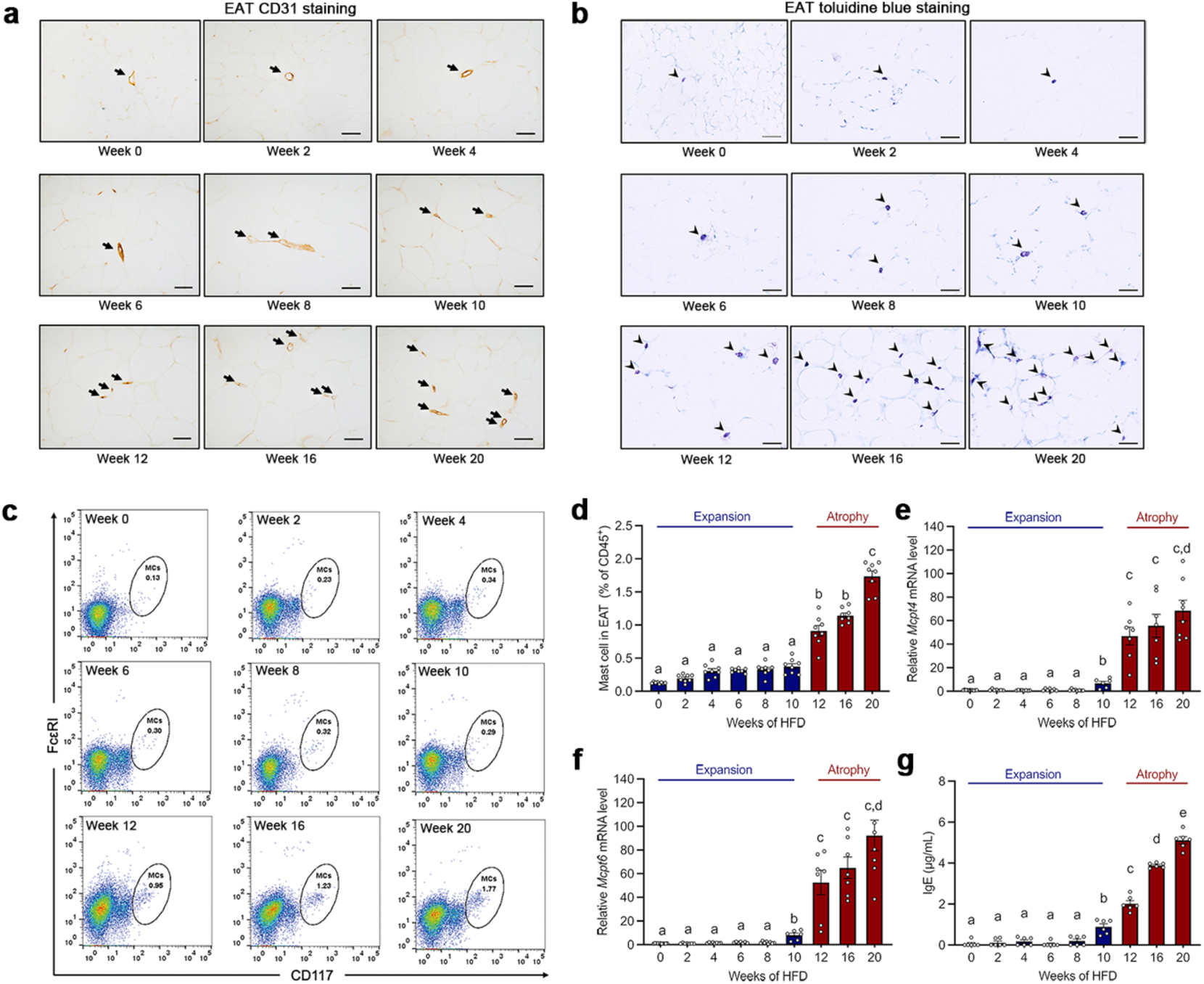
MC accumulation represents a lipolysis-associated feature during EAT atrophy. (**a/b**) Representative images of CD31 immunostaining (arrows) (**a**) and toluidine blue staining for MCs (arrowheads) (**b**) in EAT in HFD-fed WT mice. Scale bar: 50 µm. (**c/d**) Representative dot plots of flow cytometric analysis (**c**) and quantification of FcεRI^+^ CD117^+^ MCs (**d**) in EAT in HFD-fed WT mice. (**e/f**) Real-time PCR analysis of MC specific protease genes *Mcpt4* (**e**) and *Mcpt6* (**f**) in EAT in HFD-fed WT mice. (**g**) Serum IgE levels in HFD-fed WT mice. Data are mean ± SEM, n=6-8/ea. Kruskal-Wallis H test with Dunn’s post hoc adjustment for multiple testing, *p* < 0.05 was considered significant. Different letters indicate statistically significant difference.

MC accumulation and LDL-induced MC activation contribute to EAT angiogenesis in WD-fed mice^25,27^. To test whether MCs play a similar role in EAT angiogenesis in HFD-fed mice, we performed toluidine blue staining to detect MCs in EAT and SAT. The results showed progressive MC accumulation during EAT atrophy (Fig. 2b and Extended Data Fig. 5a), but MC numbers in SAT did not change (Extended Data Fig. 5b/5c). FACS analysis of EAT MCs (Fig. 2c/2d and Extended Data Fig. 5d) and EAT expression of MC specific protease genes *Mcpt4* (Fig. 2e) and *Mcpt6* (Fig. 2f) confirmed the characteristic MC accumulation. During EAT atrophy, EAT MC number correlated negatively with EAT mass and adipocyte size, but positively with liver weight, and mRNA levels of EAT *Atgl*, *Hsl*, and *Mgl*, and EAT CD31^+^ area (Extended Data Fig. 5e). Serum levels of MC activators LDL (Extended Data Fig. 3g) and IgE (Fig. 2g) were elevated during EAT atrophy, although their elevations started prior to the onset of EAT atrophy. These results suggest that MCs accumulate characteristically during EAT atrophy, and MC activation initiates adipocyte lipolysis and elevates EAT angiogenesis in HFD-fed mice.

### 3. Pharmacological stabilization of MCs ameliorates EAT atrophy and associated hepatic steatosis in HFD-fed mice

As shown in Figure 3A-1, we fed WT mice a HFD and gave daily i.p. injections of DSCG or saline. Although DSCG administration did not affect body weight gain and SAT weight (Extended Data Fig. 6a/6b), it significantly ameliorated the decrease of EAT mass and the increase of liver weight during EAT atrophy (Fig. 3b/3c). Further, we divided other HFD-fed mice into four treatment groups, including saline treatment from week 10 to 20, DSCG treatment from week 10 to 20, DSCG treatment from week 10 to 15, and DSCG treatment from week 15 to 20 (Fig. 3a-2). DSCG-treated mice exhibited comparable body weights and SAT mass to saline-treated controls (Extended Data Fig. 6c/6d). Yet, DSCG treatment from week 10 to 20 significantly reversed HFD-induced EAT mass loss and liver weight gain (Fig. 3d/3e). Notably, even DSCG treatment from week 15 to 20 was also sufficient to improve the reduction of EAT weight in HFD-fed mice (Fig. 3d).

**Figure 3.**
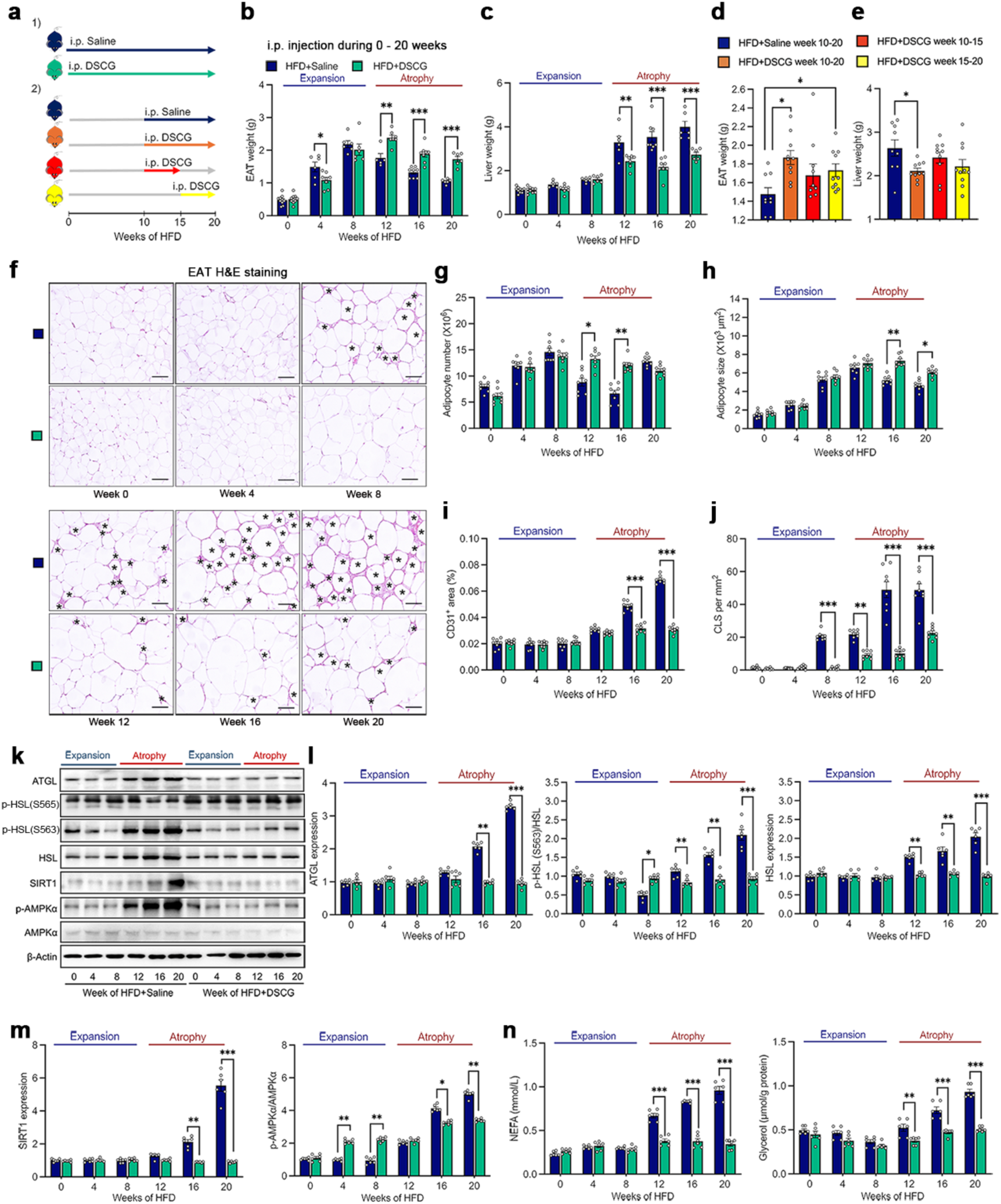
Pharmacological stabilization of MCs ameliorates EAT atrophy and associated hepatic steatosis in HFD-fed mice. (**a**) Schematic diagram of saline or DSCG treatment in HFD-fed mice. (1) WT mice received daily i.p. injections of saline (blue) or DSCG (green) from week 0 to week 20 of HFD feeding. (2) WT mice were received daily i.p. injections of saline (blue) or DSCG (orange) from week 10 to 20, daily i.p. injection of DSCG from week 10 to 15 (red), or from week 15 to 20 (yellow) of HFD feeding. (**b/c**) EAT weight (**b**) and liver weight (**c**) in WT mice received i.p. injections of saline (blue) or DSCG (green) from week 0 to 20 of HFD feeding. (**d/e**) EAT weight (**d**) and liver weight (**e**) in WT mice received i.p. injections of saline (blue) or DSCG (orange) from week 10 to 20, i.p. injections of DSCG from week 10 to week 15 (red) or from week 15 to week 20 (yellow) of HFD feeding. (**f-h**) Representative images of H&E staining (**f**), adipocyte number (**g**) and adipocyte size (**h**) in EAT in WT mice received i.p. injections of saline (blue) or DSCG (green) from week 0 to 20 of HFD feeding. Scale bars: 50 µm. Asterisks (*) indicate CLS. (**i/j**) Quantification of CD31-positive areas (**i**) and CLS number (**j**) in EAT in WT mice received i.p. injections of saline (blue) or DSCG (green) from week 0 to 20 of HFD feeding. Scale bar: 50 µm. (**k-m**) Immunoblot analysis (**k**) and quantification of (**l**) ATGL and HSL relative to β-Actin, p-HSL(563) relative to HSL, (**m**) SIRT1 and AMPKα relative to β-Actin, and p-AMPKα relative to AMPKα in EAT in WT mice received i.p. injections of saline (blue) or DSCG (green) from week 0 to 20 of HFD feeding. (**n**) Serum NEFA and glycerol levels in WT mice received i.p. injections of saline (blue) or DSCG (green) from week 0 to 20 of HFD feeding. Data are mean ± SEM, n=6-8/ea. Mann-Whitney U test for **b**, **c** and **g-n**. Kruskal-Wallis H test with Dunn’s post hoc adjustment for **d** and **e**. \**p* < 0.05, \*\**p* < 0.01, \*\*\**p* < 0.001.

We also examined the effect of DSCG treatment from week 0 to 20 on EAT adipocytes and liver phenotypes in HFD-fed mice. DSCG-treated mice had more EAT adipocytes at week 12 and 16 (Fig. 3f/3g), larger EAT adipocyte size at week 16 and 20 (Fig. 3f/3h), and lower liver lipid accumulation (Extended Data Fig. 6e/6f) than control mice during EAT atrophy. DSCG did no reduce EAT *Mcpt4* and *Mcpt6* expression or MC contents in HFD-fed mice, suggesting that DSCG inactivated MCs but did not affect MC accumulation in EAT (Extended Data Fig. 6g-6j). Relative to saline-treatment, DSCG treatment significantly reduced EAT CD31^+^ area at week 16 and 20 (Fig.3i and Extended Data Fig. 6k) and EAT CLS number from week 8 to 20 (Fig. 3j). DSCG also significantly inhibited EAT lipolysis and SIRT1/AMPK-mediated lipolytic signaling pathways (Fig. 3k-3m), and downregulated serum NEFA and glycerol levels (Fig. 3n) during EAT atrophy. Therefore, pharmacological stabilization of MCs blocks EAT adipocyte death and lipolysis, suppresses EAT angiogenesis and CLS formation, and improves EAT atrophy and associated hepatic steatosis in HFD-fed mice.

### 4. MC-derived 5-HT correlates with VAT lipolysis in HFD-fed mice and MASLD patients

Given the elevated serum levels of MC activators IgE and LDL at week 10 of HFD feeding (Fig. 2g and Extended Data Fig. 3g), we performed EAT RNA-seq analysis to detect the differential expression of genes between week 0 and 10. Nine MC mediator-related genes including *Mcpt4*, *Cma1*, *Hdc*, *Ext1*, *Ext2*, *Ndst1*, *Ndst2*, *Tph1*, and *Hpgds* were upregulated at week 10 (Fig. 4a). Real-time PCR showed that HFD feeding for 20 weeks induced EAT expression of *Mcpt4*, *Cma1*, *Hdc*, and *Tph1*, which encode MC chymases mMCP-4 and -5, histamine synthetase, and peripheral serotonin (5-HT) rate-limiting synthetase TPH1, respectively (Fig. 4b). Prior studies reported that gut-derived 5-HT elevates fasting-related adipocyte lipolysis and hepatocyte gluconeogenesis^33^. MCs rather than adipocytes or pre-adipocytes are the major source of TPH1 and 5-HT in SAT^34^. Use of ultra-high performance liquid chromatography-tandem mass spectrometry (UPLC-MS/MS) revealed that serum (Fig. 4c) and EAT (Fig. 4d/4e) 5-HT levels had increased since week 4 of HFD feeding. Relative to the baseline levels at week 0, EAT and serum 5-HT levels during EAT atrophy elevated by 5∼10-fold and 3∼4-fold, respectively (Fig. 4c-4e). DSCG treatment remarkably reduced EAT 5-HT levels from week 4 to 20 (Fig. 4e). HFD also induced the expression of EAT *Tph1*, which was suppressed completely by DSCG (Extended Data Fig. 7a/7b). To identify the sources of EAT 5-HT, we isolated adipocytes, SVFs, MCs, and MC-depleted SVFs from EAT in mice fed a HFD for 16 weeks. Similar to the expression of *Tph1* in SAT^34^, *Tph1* expression in SVFs and purified MCs from EAT was significantly higher than that in adipocytes. Deletion of MC in SVF resulted in a dramatic reduction of *Tph1* expression (Fig. 4f). Immunofluorescent double staining localized the expression of 5-HT to FcεRI^+^ and CD117^+^ MCs in EAT (Fig. 4g). Therefore, HFD-induced EAT atrophy are accompanied by increased EAT *Tph1* and 5-HT. MCs are the major source of TPH1 and 5-HT in EAT in HFD-fed mice. Therefore, elevated serum LDL and IgE during EAT atrophy may activate EAT MCs to secrete 5-HT and promote EAT lipolysis and NEFA release. This hypothesis is supported by the observation that there were a series of positive correlations in serum MC-relevant markers, including those between LDL and 5-HT, IgE and 5-HT, LDL and NEFA, IgE and NEFA, and 5-HT and NEFA in HFD-fed mice (Fig. 4h). Except for the correlation between LDL and 5-HT levels, other serum marker correlations were also observed in patients clinically diagnosed with MASLD (Fig. 4i). Further, such correlations were independent of whether the MASLD patients possessed metabolic associated steatohepatitis (MASH) or not (Extended Data Fig. 8a/8b).

**Figure 4.**
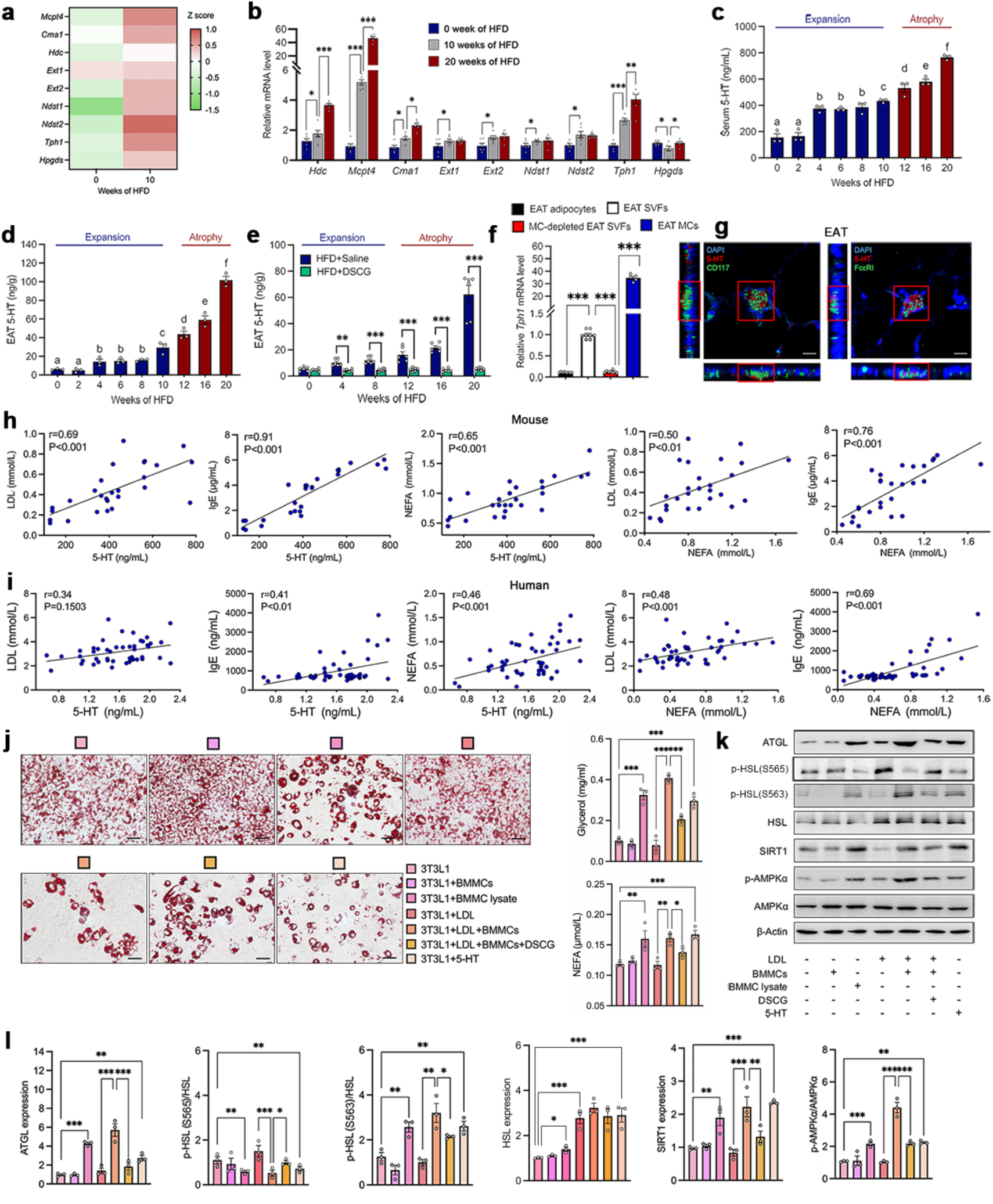
MC-derived 5-HT is correlated with VAT lipolysis in HFD-fed mice and MASLD patients, and promotes adipocyte lipolysis. (**a**) Heatmap of MC mediator-related gene expression in EAT in WT mice at week 0 and at week 10 of HFD feeding. n=3/ea. (**b**) Real-time PCR analysis of MC mediator-related genes *Mcpt4* (encodes mouse mast cell protease 4 (mMCP-4), *Cma1* (encodes mMCP-5), *Hdc* (encodes histidine decarboxylase), *Ext1* (encodes exostosin-1), *Ext2*, *Ndst1* (encodes N-deacetylase/N-sulphotransferase 1), *Ndst2*, *Tph1* (encodes tryptophan hydroxylase 1), and *Hpgds* (encodes hematopoietic prostaglandin D synthase) in EAT in WT mice at week 0, 10, 20 of HFD feeding. n=6-8/ea. (**c/d**) Serum (**c**) and EAT (**d**) 5-HT levels in HFD-fed WT mice. n=6-8/ea. (**e**) EAT 5-HT levels in WT mice received i.p. injections of saline or DSCG from week 0 to 20 of HFD feeding. n=6-8/ea. (**f**) Real-time PCR analysis of *Tph1* mRNA levels in adipocytes, SVFs, MC-depleted SVFs, and MCs from WT EAT in HFD-fed mice for 16 weeks. n=5-8/ea. (**g**) 5-HT^+^ (red) and CD117^+^ (green), 5-HT^+^ (red) and FceRI^+^ (green) immunofluorescent double staining in WT EAT in HFD-fed mice for 16 weeks. (**h**) Correlations between serum LDL and 5-HT levels, IgE and 5-HT levels, LDL and NEFA levels, IgE and NEFA levels, and 5-HT and NEFA levels during EAT atrophy in HFD-fed mice. n=27. (**i**) Correlations between serum LDL and 5-HT levels, IgE and 5-HT levels, LDL and NEFA levels, IgE and NEFA levels, and 5-HT and NEFA levels in MASLD patients. n=50. (**j**-**l**) Differentiated 3T3-L1 adipocytes were treated with or without live BMMCs, BMMC lysates, LDL, LDL-activated BMMCs, DSCG- and LDL-treated BMMCs, or 5-HT. n=3/ea. (**j**) Representative images of oil-red O staining, media glycerol and NEFA levels in 3T3-L1 adipocytes. Scale bars: 50 µm. (**k/l**) Immunoblot analysis (**k**) and quantification of (**l**) ATGL, HSL and SIRT1 relative to β-actin, p-HSL(565) and p-HSL(563) relative to HSL, and p-AMPKα relative to AMPKα in 3T3-L1 adipocytes. Data are mean ± SEM. Pearson’s correlation test for **h** and **i**. P < 0.05 was considered significant correlation. Mann-Whitney U test for **e**. Kruskal-Wallis H test with Dunn’s post hoc adjustment for **b-d**, and **f**. Welch’s t-test for **j** and **l**. \**p* < 0.05, \*\**p* < 0.01, \*\*\**p* < 0.001. Different letters indicate statistically significant difference.

### 5. MC-derived 5-HT promotes adipocyte lipolysis

MCs are known to express *Tph1* and secrete 5-HT after stimulation with LDL^27,34^. Here we performed a linear regression analysis and showed that EAT 5-HT level correlated positively with serum LDL during EAT atrophy (Fig. 4h/4i). To test whether MC-derived 5-HT drove adipocyte lipolysis, we isolated EAT explants from MC-deficient *Kit^w-sh/w-sh^*mice. We treated these EAT explants with bone marrow-derived MC (BMMC) lysates and physiological concentration of 5-HT (100 ng/mL). The results showed that both BMMC lysates and 5-HT promoted the releases of glycerol and NEFA and enhanced the expression of *Atgl* in these EAT explants (Extended Data Fig. 8c). Further, in differentiated 3T3-L1 adipocytes, BMMC lysates, LDL-activated BMMCs, and 5-HT, but not live BMMCs or LDL alone reduced adipocyte lipid storage and elevated glycerol and NEFA releases (Fig. 4j and Extended Data Fig. 8d). Along with the promotion of lipolysis, BMMC lysates, LDL-activated BMMCs, and 5-HT also induced the expression and activation of lipolytic enzymes and SIRT1/AMPK-mediated lipolytic signaling pathways (Fig. 4k/4l and Extended Data Fig. 8e). DSCG treatment reversed the effects of LDL-activated BMMCs on lipolytic enzymes and SIRT1/AMPK signaling molecules in differentiated 3T3-L1 adipocytes (Fig. 4k/4l and Extended Data Fig. 8e). These observations demonstrate that the 5-HT release from activated MCs promotes adipocyte lipolysis *via* the SIRT1 and AMPKα signaling pathways.

### 6. MC-specific depletion of *Tph1* ameliorates EAT atrophy and hepatic steatosis in HFD-fed mice

Further, we selectively depleted *Tph1* in MCs by crossbreeding *Mcpt4-Cre* mice with *Tph1^fl/fl^* (Tph1-Con) mice to generate *Mcpt4-Cre; Tph1^fl/fl^* (Tph1-MCKO) mice (Fig. 5a). Compared to the Tph1-Con mice, Tph1-MCKO mice exhibited significantly reduced MC *Tph1* expression in SAT, EAT and peritoneal cavity (Extended Data Fig. 9a) and the 5-HT levels in SAT and EAT (Extended Data Fig. 9b). MC-selective depletion of *Tph1* did not affect mouse body weight gain after CD or HFD feeding (Extended Data Fig. 9c). Like those in the WT C57BL/6N mice, HFD also induced an EAT remodeling with initial EAT expansion and subsequent EAT atrophy in Tph1-Con mice (Fig. 5b). During EAT atrophy, HFD-fed Tph1-Con mice showed increased liver weight and lipid deposition (Fig. 5c and Extended Data Fig. 9d), declined and then restored adipocyte number (Fig. 5e/5f), reduced adipocyte size (Fig. 5e/5g), increased and then decreased CLS number (Extended Data Fig. 9e), elevated angiogenic CD31^+^ area (Extended Data Fig. 9f), and activation of lipolytic enzymes and associated AMPK/SIRT1 signaling molecules (Fig. 5h-5j). MC-specific depletion of *Tph1* reversed those aforementioned HFD-induced changes (Fig. 5b-5j and Extended Data Fig. 9d-9f). These observations support the hypothesis that MC-derived 5-HT not only leads to EAT atrophy and hepatic steatosis but also regulates EAT adipocyte turnover and angiogenesis by promoting adipocyte lipolysis.

**Figure 5.**
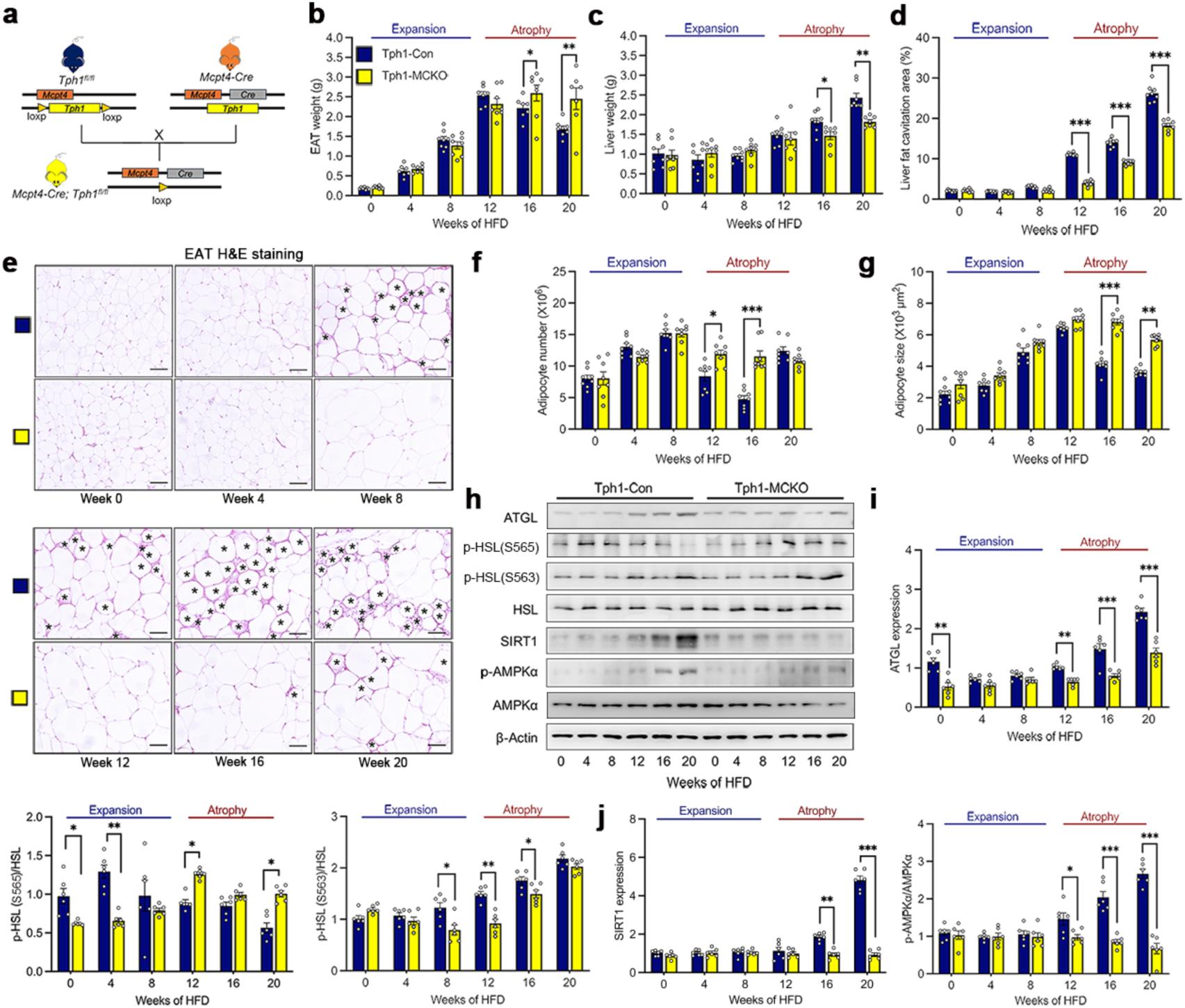
MC-specific depletion of *Tph1* improves EAT atrophy and hepatic steatosis in HFD-fed mice. (**a**) *Tph1^fl/fl^* mice were crossbreed with *Mcpt4-Cre* mice to generate *Mcpt4-Cre; Tph1^fl/fl^* (Tph1-MCKO) mice and littermate *Tph1^fl/fl^* (Tph1-Con) mice. (**b/c**) EAT weight (**b**) and liver weight (**c**) in Tph1-Con and Tph1-MCKO mice during HFD feeding for 20 weeks. (**d**) Percentage of liver fat cavitation area in Tph1-Con and Tph1-MCKO mice during HFD feeding for 20 weeks. (**e-g**) Representative images of H&E staining (**e**), adipocyte number (**f**) and adipocyte size (**g**) in EAT in Tph1-Con and Tph1-MCKO mice during HFD feeding for 20 weeks. Scale bars: 50 µm. Asterisks (*) indicate CLS. (**h-j**) Immunoblot analysis (**h**) and quantification of (**i**) ATGL relative to β-Actin, p-HSL(565) and p-HSL(563) relative to HSL, (**j**) SIRT1 relative to β-Actin, and p-AMPKα relative to AMPKα in EAT in in Tph1-Con and Tph1-MCKO mice during HFD feeding for 20 weeks. Data are mean ± SEM, n=6-8. Mann-Whitney U test. \**p* < 0.05, \*\**p* < 0.01, \*\*\**p* < 0.001.

### 7. Adipocyte-selective ablation of *Htr2b* reverses EAT atrophy and hepatic steatosis in HFD-fed mice

5-HT exerts its biological functions after binding to multiple 5-HT receptors (HTRs)^35^. To identify the exact HTR that mediates the functions of MC-derived 5-HT in promoting adipocyte lipolysis, we tested the expression of several HTR genes in EAT from WT mice fed a HFD for 10 and 20 weeks. Among these HTRs, only *Htr2b* was upregulated in EAT after 10 and 20 weeks of HFD feedings (Fig. 6a). When 3T3-L1 adipocytes were treated with BMMC lysates and 5-HT to increase lipolysis as illustrated by reduced intracellular oil-red O staining (Extended Data Fig. 10a), the expression of *Htr2b* showed most robust increase among all tested HTRs (Fig. 6b). In HFD-fed mice, along with the increased expression of EAT *Tph1* (Extended Data Fig. 7), EAT *Htr2b* expression showed concurrent increase (Fig. 6c/6d). DSCG administration suppressed *Htr2b* expression during EAT atrophy (Fig. 6d). To test whether adipocyte *Htr2b* mediates the effects of MC 5-HT on EAT atrophy and associated hepatic steatosis in HFD-fed mice, we selectively depleted *Htr2b* in adipocytes by crossbreeding *Adipoq-Cre* mice with *Htr2b^fl/fl^* (Htr2b-Con) mice to produce *Adipoq-Cre; Htr2b^fl/fl^* (Htr2b-AKO) mice (Fig. 6e). Compared to Htr2b-Con mice, Htr2b-AKO mice showed successful depletion of EAT *Htr2b* expression without affecting *Htr2a* expression (Extended Data Fig. 10b), MC number (Extended Data Fig. 10c/10d), or *Tph1* expression (Extended Data Fig. 10e) in EAT. Adipocyte-selective depletion of *Htr2b* did not affect body weight gain in HFD-fed mice (Extended Data Fig. 10f), but significantly resisted HFD-induced EAT atrophy and hepatic steatosis (Fig. 6f-6h and Extended Data Fig. 10g). Relative to Htr2b-Con mice, Htr2b-AKO mice showed more EAT adipocyte number at week 16 (Fig. 6i/7j), larger EAT adipocyte size at week 16 and 20 (Fig. 6i/6k), fewer EAT CLS numbers from week 8 to 20 (Extended Data Fig. 10h), smaller EAT CD31^+^ area at week 16 and 20 (Fig. 6i and Extended Data Fig. 10j), and lower levels of lipolytic enzymes and AMPK/SIRT1 signaling molecules (Fig. 6l-6n) during EAT atrophy. Together, our observations from HFD-fed mice suggest that MC-driven adipocyte lipolysis leads to EAT atrophy and hepatic steatosis, a mechanism involving interactions between MC-derived 5-HT and its receptor HTR2b on adipocytes.

**Figure 6.**
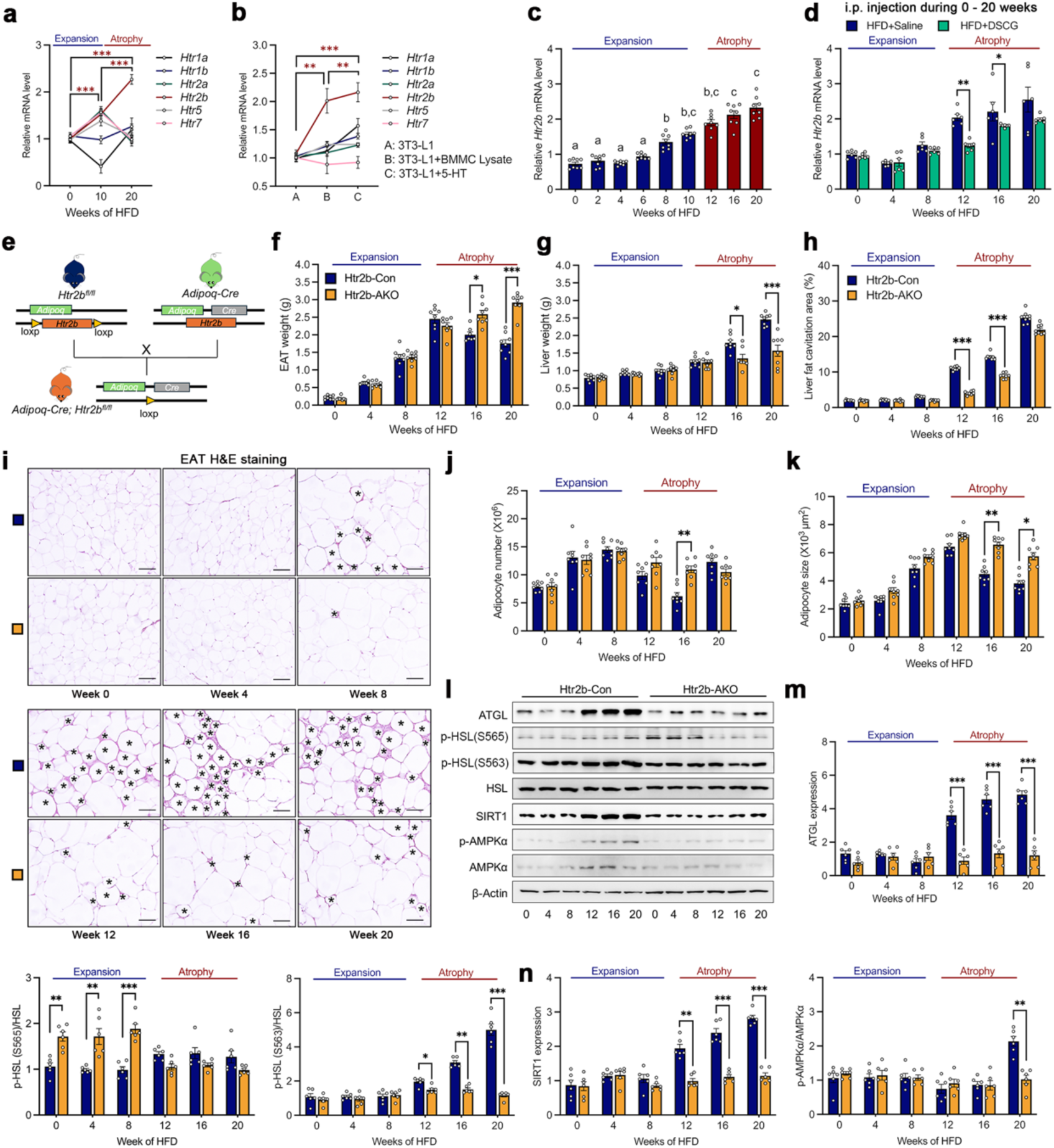
Adipocyte-selective ablation of *Htr2b* reverses EAT atrophy and hepatic steatosis in HFD-fed mice. (**a**) Real-time PCR analysis of 5-HT receptor (Htr) genes *Htr1a*, *Htr1b*, *Htr2a*, *Htr2b*, *Htr5* and *Htr7* mRNA levels in EAT in WT mice at week 0, 10, and 20 of HFD feeding. n=6-8/ea. (**b**) Real-time PCR analysis of *Htr1a*, *Htr1b*, *Htr2a*, *Htr2b*, *Htr5* and *Htr7* expression in differentiated 3T3L1 adipocytes treated with or without BMMC lysates and 5-HT. n=3/ea. (**c**) EAT *Htr2b* mRNA levels in WT mice during HFD feeding for 20 weeks. n=6-8/ea. (**d**) EAT *Htr2b* mRNA levels in WT mice received i.p. injections of saline or DSCG from week 0 to 20 of HFD feeding. n=6-8/ea. (**e**) *Htr2b^fl/fl^* mice were crossbreed with *Adipoq-Cre* mice to generate *Adipoq-Cre; Htr2b^fl/fl^* (Htr2b-AKO), *Htr2b^fl/fl^* (Htr2b-Con) littermate mice. (**f/g**) EAT weight (**f**) and liver weight (**g**) in Htr2b-Con and Htr2b-AKO mice during HFD feeding for 20 weeks. (**h**) Percentage of liver fat cavitation area in Htr2b-Con and Htr2b-AKO mice during HFD feeding for 20 weeks. n=6-8/ea. (**i-k**) Representative images of H&E staining (**i**), adipocyte number (**j**) and adipocyte size (**k**) in EAT in Htr2b-Con and Htr2b-AKO mice during HFD feeding for 20 weeks. Scale bars: 50 µm. The asterisks (*) indicates the CLS. n=6-8/ea. (**l-n**) Immunoblot analysis (**l**) and quantification of (**m**) ATGL relative to β-Actin, p-HSL(565) and p-HSL(563) relative to HSL, (**n**) SIRT1 relative to β-Actin, and p-AMPKα relative to AMPKα in EAT in in Htr2b-Con and Htr2b-AKO mice during HFD feeding for 20 weeks. n=6-8/ea. Data are mean ± SEM. Kruskal-Wallis H test with Dunn’s post hoc adjustment for **c**. Mann-Whitney U test for **d, f-h, j, k, m and n**. \**p* < 0.05, \*\**p* < 0.01, \*\*\**p* < 0.001. Different letters indicate statistically significant difference.

## Discussion

This study reports a previously untested function of MCs and an unanticipated mode of communication between MCs and adipocytes (Fig. 7), originated from our initial observations that MC accumulation in EAT represents a lipolysis-associated feature in HFD-fed mice. Adipocyte lipolysis is the primary driver of post-expansive EAT lipoatrophy. Pharmacological stabilization of MCs reduces EAT lipolysis and ameliorates EAT atrophy and hepatic steatosis. Activated MCs directly promote adipocyte lipolysis by releasing 5-HT. We employed both MC-specific depletion of *Tph1* and adipocyte-selective ablation of *Htr2b* in HFD-fed mice to demonstrate that MC-derived 5-HT binds to adipocyte HTR2b, thereby triggering adipocyte lipolysis and subsequently inducing EAT atrophy and hepatic steatosis.

**Figure 7.**
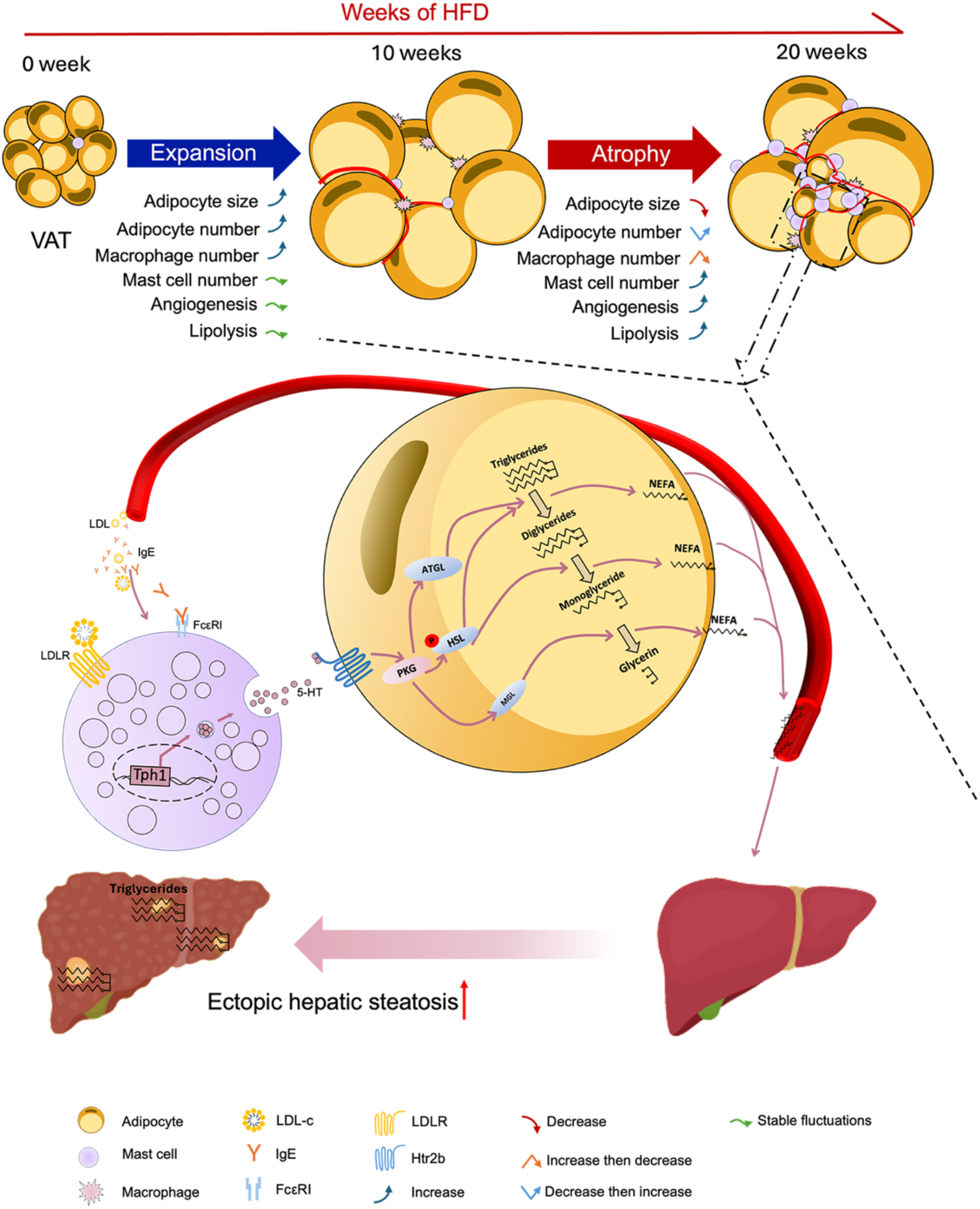
Proposed mechanisms of MC-driven adipocyte lipolysis leading to VAT atrophy and ectopic hepatic steatosis in HFD-fed mice. HFD feeding induces EAT remodeling including “expansion” from week 0 to 10 and “atrophy” from week 12 to 20. In the atrophic EAT, adipocyte size reduction is accompanied by increased angiogenesis, lipolysis, and MC accumulation. Meanwhile, serum levels of MC activators LDL and IgE are progressively elevated to activate MCs to release 5-HT through LDLR and FcεR1. MC-derived 5-HT triggers adipocyte lipolysis and further leads to EAT atrophy and hepatic steatosis through HTR2b-mediated SIRT1/AMPKα signaling.

As a highly plastic and dynamic organ, WAT has a remarkable ability to expand its mass and volume in response to excess nutrients *via* adipocyte hypertrophy and/or hyperplasia. In HFD-fed mice, anatomical localization of WAT into subcutaneous or visceral depots displayed distinct cellular alterations accompanying adipose tissue expansion and linked to obesity-associated hepatic metabolic risks^8,9^. In mice under a CD, SAT possessed smaller adipocyte size, slower triglyceride turn over, lower oxidative stress, and higher adipocyte number and respiratory capacity than those of EAT^36,37^. After HFD feeding however, SAT expanded predominantly through adipocyte hypertrophy, whereas EAT expansion involved both adipocyte hyperplasia and hypertrophy. EAT adipocyte hyperplasia was initiated after 4 weeks of HFD^38-40^. In contrast, SAT maintained a similar degree of vascularization to that under CD conditions^41^, suggesting continuous adipocyte hypertrophy and SAT expansion without tissue inflammation from hypoxia and nutrient deficiency^8,42^. In EAT from HFD-fed mice, vascular structures were reduced compared to those in EAT from CD-fed mice. M1 macrophages are one of the most common pro-inflammatory cells that were recruited into EAT where these macrophages formed CLS^8,42^. Along with insufficient angiogenesis and progressive inflammation in EAT from HFD-fed mice, prolonged HFD consumption led EAT adipocytes to reach to their limits of lipid storage capacity^43^, and even cause EAT adipocyte size and tissue mass reductions^8^, resulting in ectopic fat deposition into the liver. Therefore, in HFD-fed mice, unlike the continuous expansion of SAT, EAT displayed a characteristic transition of expansion-to-atrophy, reflecting the fundamental metabolic heterogeneity between these different white adipose depots. In agreement with these prior observations, our results demonstrate that adipocyte size gradually enlarged during EAT expansion in HFD-fed mice (Fig. 1e/1g), and adipocyte hyperplasia (Fig. 1e/1g) and macrophage infiltration (Extended Data Fig. 4c-4f) occurred after 4 weeks of HFD. When mice entered to the post-expansive EAT atrophy stage, hypertrophic adipocytes showed sustained decrease in size (Fig. 1e/1g). The inverse relationship between adipocyte and macrophage numbers (Fig. 1f and Extended Data Fig. 4b-4f) supports that macrophage-associated adipocyte death and CLS formation peaked at week 12 and 16 with subsequent resolution at week 20 (Extended Data Fig. 2c-2f). MC accumulation (Fig. 2b-2f and Extended Data Fig. 5a), lipolysis (Fig. 1h-1j and Extended Data Fig. 3b/3c), and angiogenesis (Fig. 2a and Extended Data Fig. 4a) in EAT as well as hepatic steatosis (Fig. 1d and Extended Data Fig. 1g-1i) were characteristically elevated during EAT atrophy. Pharmacological stabilization of MCs nearly counteracted all these alterations in atrophic EAT, spanning adipocytes, ECs, and macrophages, and even extending hepatocytes through EAT NEFA release (Fig. 3 and Extended Data Fig. 6). Therefore, MC accumulation and activation in EAT play a pivotal role in HFD-induced EAT atrophy.

MCs are tissue-resident sentinel cells possessing much broader than their original role in IgE-mediated allergic disease^23,24^. Mouse MCs are subclassified into two subtypes, connective tissue MCs (CTMCs) that express mouse MC proteases mMCP-4, -5, -6, and carboxypeptidase A and mucosal MCs (MMCs) that preferentially express mMCP-1 and -2^43,44^. In this study, we reported upregulation of *Mcpt4*, *Cma1*, and *Mcpt6* that encode CTMC-specific proteases mMCP-4, -5, and -6 in EAT MCs during EAT atrophy (Fig. 2b-2f/4a/4b). Therefore, we used *Mcpt4-Cre* mice to generate MC-selective TPH1-deficient mice to examine the role of MC-derived 5-HT in EAT adipocyte lipolysis and HFD-induced EAT atrophy (Fig. 5 and Extended Data Fig. 9). In mammals, the central and peripheral 5-HT is primarily synthesized by the rate-limiting enzymes TPH2 and TPH1, respectively^45^. In mice, fasting-induced increases in circulating 5-HT, NEFA, and glycerol levels are accompanied by reductions of EAT weight and adipocyte size. Depletion of either gut *Tph1* or adipocyte *Htr2b* blunted the effects of fasting, suggesting that gut-derived 5-HT initiated EAT lipolysis *via* adipocyte HTR2b signaling^33^. In mice fed a HFD for 8-10 weeks, WAT expansion is accompanied with enhanced expression of *Tph1* and 5-HT in EAT and SAT, but the expression of *Htr2a* and *Htr2b* was elevated only in EAT^46,47^. Inhibition and depletion of adipocyte *Htr2b* reduced EAT lipolysis and improved systemic insulin resistance^46^. In contrast, adipocyte 5-HT synthesis enhanced EAT lipogenesis *via* HTR2a signaling^47^. In mice fed a HFD for 12 weeks, inhibition of gut-derived 5-HT synthesis reduced circulating NEFA and glycerol^33^. Therefore, under different dietary conditions, peripheral 5-HT from distinct cell sources in gut and adipose tissue fine-turns the balance and dynamics of lipogenesis and lipolysis in EAT adipocytes through HTR2a and HTR2b signaling pathways, respectively.

In above studies of fasting-induced EAT lipolysis and HFD-induced EAT expansion, adipocyte-specific ablation of *Htr2b* affected only HSL phosphorylation without altering ATGL level in EAT^33,46^, implying that 5-HT does not trigger large-scale EAT lipolysis under these dietary conditions. In mice fed a HFD for 12 weeks, depletion of gut *Tph1* and pharmacological restoration of circulating 5-HT to normal level both reduced plasma NEFA and glycerol, but failed to normalize plasma NEFA and/or glycerol levels to those in CD-fed WT mice^33^. These observations suggest that non-gut-derived 5-HT also controls EAT lipolysis during HFD-induced EAT atrophy. Here we report that MCs are the primary source of 5-HT in EAT. MC activation and physiological 5-HT concentration are sufficient to trigger adipocyte lipolysis (Fig. 4j-4l and Extended Data Fig. 8c-8e). Although both *Htr2a* and *Htr2b* were increased during EAT expansion, EAT atrophy is only accompanied with persistent elevation of *Htr2b* (Fig. 6a). MC-specific *Tph1* deletion (Fig. 5) or adipocyte *Htr2b* ablation (Fig. 6) repressed the elevation of ATGL expression and HSL activity during EAT atrophy. Circulating MC activators (IgE and LDL) were elevated during EAT expansion (Fig. 2g and Extended Data Fig. 3g). At week 8 of HFD feeding, HSL phosphorylation in EAT was reduced following MC inactivation or MC-selective *Tph1* deletion (Fig. 3k/3l/5h/5i), supporting a role for MC-derived 5-HT in adipocyte lipolysis during EAT expansion. Together, in HFD-induced transition from EAT expansion to atrophy, gut-and MC-derived 5-HT may collectively mediate modest lipolysis during EAT expansion, whereas MC-derived 5-HT plays an indispensable role in driving massive lipolysis during EAT atrophy.

In this study, we report that MC-driven lipolysis represents a core biological event in metabolic abnormities during EAT atrophy. Similar to our results, reduced lipolysis from genetic deficiency of lipolytic enzymes ATGL and HSL or adipocyte-specific expression of G0S2 led to a metabolically healthy obesity phenotype, whereas enhanced basal lipolysis contributed to clinical pathogenesis of obesity-associated metabolic abnormalities^48^. In HFD-fed mice, adipocyte size increase peaked at the onset of EAT atrophy and progressively declined throughout the EAT atrophy^8^. Macrophage infiltration, CLS formation, and adipocyte death peaked during EAT atrophy, suggesting that EAT adipocyte lipolysis facilitates macrophage-associated adipocyte death and CLS formation^8^. Loss of endothelial *Pten* elevated WAT angiogenesis and stimulates adipocyte lipolysis *via* polyamine production and EC proliferation that relies on the presence of fatty acids^49^. In fasted mice, adipocyte-originated prostaglandin E2 (PGE2) mediated WAT macrophage accumulation during lipolysis^50^. Adipocyte PGE2-EP4 axis promoted WAT lipolysis to induce ectopic hepatic steatosis^48^. Absence of adipocyte ATGL impaired fasting-induced liver ketogenesis and FGF21 production by controlling hepatocyte PPARα activity^51^. Interestingly, fasting stimulated extra-hepatic MCs to release histamine to trigger liver production of endogenous PPARα agonist oleoylethanolamide (OEA). Together with NEFAs, OEA activated hepatic *Pparα* and associated ketogenesis, whereas HFD dampened MC histamine release and liver OEA production^52^. MC mediators 5-HT and histamine may modulate HFD-induced metabolic abnormalities in EAT and liver, including adipocyte death, lipolytic activation, and angiogenesis elevation in EAT, hepatic steatosis and *Pparα* downregulation during EAT atrophy, a hypothesis merits further investigation.

As we summarized in Fig. 7, HFD induced two phases of EAT remodeling, starting with EAT expansion followed by EAT atrophy. An unappreciated communication mode between MCs and adipocytes drives large-scale lipolysis to lead to post-expansive VAT atrophy and ectopic hepatic steatosis. Positive correlations among circulating molecules, including IgE, LDL, 5-HT, and NEFA, are identified in HFD-fed mice and MASLD patients. Given the increasing global burden of obesity and MASLD and no approved pharmacological agents for MASLD^3,53,54^, cellular communication of MCs and adipocytes *via* the interaction between 5-HT and HTR2b shows promise as a therapeutic target for obesity-associated MASLD.

## Methods

### Mice

Wild-type (C57BL/6N) and *Kit^w-sh/w-sh^* (C57BL/6J) mice were purchased from Vital River Laboratory Animal Technology Co. Ltd. (Beijing, China) and the Jackson Laboratories (Bar Harbor, ME, USA), respectively. *Tph1^fl/fl^* (C57BL/6N) and *Htr2b^fl/fl^* (C57BL/6N) mice were obtained from GemPharmatech LLC. (Jiangsu, China). *Mcpt4-Cre* (C57BL/6N) and *Adipoq-Cre* (C57BL/6J) mice were from Cyagen Biosciences Inc. (Jiangsu, China). Mice were housed in ventilated cages within a pathogen free barrier facility maintained on a 12-hr light and 12-hr dark cycle. All experimental procedures were approved by the Animal Committee of Hefei University of technology. To induce DIO, 5-week-old WT mice, Tph1-MCKO mice, Tph1-Con mice, Htr2b-AKO mice and Htr2b-Con mice were fed on an HFD (45% calories from fat, Research Diet Cat # D12451 or Jiangsu Xietong Medicine Bioengineering Co. Cat# XTM04-002), and age-matched WT mice for a CD (Jiangsu Xietong Medicine Bioengineering Co. Cat# XTI01SL-002). To stabilize MCs, WT HFD-fed mice were given daily i.p. injections of DSCG (25 mg/kg body weight/day) or saline. Animal body weight was monitored weekly.

### Histology

Adipose tissues and liver were fixed in 4% formalin, embedded in paraffin, and serially sliced into 6 µm thickness. For histological characterizations of adipose tissue and liver, hematoxylin & eosin (H&E) staining were performed. EAT apoptotic cells were determined by TUNEL FITC Apoptosis Detection Kit (Yeasen Biotech, Shanghai, China Cat# 40306ES), according to the manufacturer’s instructions. For immunohistochemical analysis of macrophage and angiogenesis, mouse EAT sections were stained with rat anti-Mac2 (1:800, Bio Legend, San Diego, CA, USA Cat# ab56297) and rabbit anti-CD31 (1:300, Abcam, Cambridge, MA, USA Cat# ab28364) antibodies, followed by appropriate biotin-conjugated secondary antibodies and HRP-streptavidin. Using the Image-Pro Plus software (Media Cybernetics, Bethesda, MD), we quantified adipocyte size, liver fat cavitation, CLS number, numbers of TUNEL-positive cells, Mac2- and CD31-positive areas in five random fields in each stained section.

MC-specific toluidine blue staining was performed and MC number in adipose tissues was quantified as previous reported^27^. For immunofluorescent co-localization analyses, EAT sequential sections were stained with mouse anti-5-HT (1:200, NovusBio, Shanghai, China Cat# NB120-16007), rabbit anti-CD117 (1:200, Abcam, Cambridge, MA, USA Cat# 135108), and rabbit anti-FcεR1 (1:200, Abcam, Cambridge, MA, USA Cat# ab288427) antibodies, followed by fluorophore-conjugated secondary antibody (550-conjugated antibody for 5-HT detection, 488-conjugated antibody for CD117 or FcεR1 detection). Sections were counterstained with DAPI (422801, Biolegend Cat# 422801), and Z-stack scanning of images were captured with a confocal microscopy (ZEISS, Germany).

### Real-time PCR

Total RNA was extracted from tissues or cells using Trizol reagent (Takara, Dalian, China Cat# 1725150) and reverse-transcribed into cDNA using a reverse transcription kit (Takara) according to the manufacturer’s instructions. Quantitative real-time PCR was performed by iTaq universal SYBR Green supermax (Bio-Rad) in Bio-Rad MyiQ2 real-time PCR system. The relative mRNA expression was detected by adopting ΔΔCT method and normalized to β-actin. The primer sequences were listed in Table S2.

### Immunoblot analysis

For immunoblot analysis, an equal amount of proteins extracted from EAT were separated on SDS-PAGE and immunoblotted with the following primary antibodies including rabbit anti-ATGL (1:1000, Cell Signaling Technology, Danvers Cat# 2138S), rabbit anti-HSL (1:500, Abcam Cat# 4107S), rabbit anti-p-HSL(S565) (1:500, Abcam Cat# ab109400), rabbit anti-p-HSL(S563) (1:500, Abcam Cat# ab62153), rabbit anti-SIRT1 (1:1000, Cell Signaling Technology Cat# 2028S), rabbit anti-AMPKα (1:1000, Cell Signaling Technology Cat# 2532S), rabbit anti-p-AMPKα (1:1000, Cell Signaling Technology Cat# 2535S), rabbit anti-Caspase-3 (1:1000, Cell Signaling Technology Cat# 9661), rabbit anti-RIP3 (1:1000, Cell Signaling Technology Cat# 95702), rabbit anti-p-RIP3 (1:1000, Cell Signaling Technology Cat# 93654), and rabbit anti-β-actin (1:1000, Cell Signaling Technology Cat# 8457S), followed by appropriate HRP-conjugated secondary antibodies. Chemiluminescence was detected by ECL western blot kit (ThermoFisher Cat# 34580) in Image Quant LAS 4000 mini (GE Healthcare, Piscataway, NJ, USA).

### Flow cytometry

SVFs were isolated by collagenase digestion from EAT, and re-suspended in FACS buffer after red blood cell lysis. After the incubation with an Fc block (BioLegend), SVFs cells were stained with PE anti-mouse CD45 (BioLegend Cat# 103106), PE-Cyanine7 anti-mouse FceR1 (BioLegend Cat# 25-5898-82), APC anti-mouse CD117 (BioLegend Cat# 135108) for 30 min at 4 °C. Cells were initially selected by size on the basis of forward scatter (FSC) and side scatter (SSC), and then sorted on the basis of cell-surface markers using a Beckman MoFlo XDP (Beckman Coulter, CA, USA),

### Serum and tissue biochemistry

Serum levels of TC, TG, HDL, and LDL were detected using a biochemical analyzer (Beckman Coulter, AU680). Glycerol levels were detected using a glycerol detection kit (Sigma-Aldrich, Cat# F6428), and NEFA levels were detected using a free fatty acid kit (Fujifilm, Cat# 294-63601) according to the manufacturer’s instructions. EAT 5-HT levels were determined using UPLC-MS/MS (Thermo Fisher Scientific, Accela 600 pump UPLC system equipped with LTQ Orbitrap XL mass spectrometer) as described previously^34^. Serum 5-HT (LDN, Nordhorn, Germany Cat# BA E-5900R) and IgE (JYM ELISA Lab, Wuhan, China Cat# JYM1391Mo) levels were detected using mouse ELISA kits according to manufacturer protocols.

### MASLD patients

We obtained serum samples from MASLD patients from the Affiliated Hospital of Wenzhou Medical University. A total of 50 adult MASLD patients, including 25 MASH patients and 25 non-MASH patients, participated in this study. Detailed human donor demographic information is provided in Table S1. The study protocol was approved by the ethics committee of the First Affiliated Hospital of Wenzhou Medical University. Written informed consent was obtained from each participant.

### Cell culture

BMMCs were induced from bone marrow from 8-week-old mice as we reported previously^25,27,34^. After 5 weeks, cell purity was confirmed by flow cytometry to show the cell surface expression of FcεRI and CD117 (> 98%). To activate BMMCs, the cells were treated with 1 mg/mL LDL (Lee Biosolutions, Inc, Maryland Heights, MO Cat# 360-10). LDL-treated BMMCs were inactivated with 100 nM DSCG.

For 3T3-L1 adipocyte differentiation, confluent cells were cultured in DMEM supplemented with 10% FBS, 10 mg/mL insulin, 0.5 mM isobutylmethylxanthine, and 1 mM dexamethasone (all from Sigma-Aldrich) for 2 days, and then maintained in DMEM containing insulin for another 10 days. After differentiation, adipocytes were treated with live BMMCs (1×10^7^ cells per well for 12-well plate), BMMC lysate (equivalent to the live cell numbers), LDL (1 mg/mL), or 5-HT (100 ng/mL) for 48 hrs. To assess adipocyte neutral lipid levels, cells were stained with oil-red O (Sigma Cat# O0625) and quantified lipid accumulation by absorbance at OD510 nm.

### Statistical analysis

All data were expressed as mean ± SEM. For the correlation analysis, Pearson’s correlation test was used. Non-normally distributed murine outcomes were compared between two independent groups using the Mann-Whitney U test, while multi-group comparisons were analyzed via the Kruskal-Wallis H test with Dunn’s post hoc adjustment for multiple testing. For cell experiments, Welch’s t-test was employed for group comparisons after confirming approximate normality. \**p* < 0.05, \*\**p* < 0.01, \*\*\**p* < 0.001.

## Data availability

Mouse lines and cell lines generated in this study will be available through the lead contact upon reasonable request. The published article includes all datasets generated or analyzed during this study and available from the lead contact upon request. Further information and requests for resources and reagents should be directed to Jian Liu.

## Acknowledgements

This work is supported by the National Natural Science Foundation of China (32070757 to JL and 32200630 to XZ) and the Fundamental Research Funds for the Central Universities (PA2025GDGP0026 to JL and JZ2024HGTB0242 to XZ).

## Author contributions

J.W. and L.Z. designed and performed experiments, analyzed data, generated the figure panels, and wrote the draft manuscript. M. Z., X.J., W.L., H.Y., W.Y. and W.H. provided the blood samples of human. H.Y., H.C., and B.B. helped to perform the experiments. G.P. helped to design experiments and revised the manuscript. X.Z. and J.L. lead the project, designed the experiments, and revised the manuscript.

