## Supplementary data for "Mast cell-driven adipocyte lipolysis promotes post-expansive VAT atrophy and ectopic hepatic steatosis"

1. School of Food and Biological Engineering, Hefei University of Technology, Hefei 230009, China
2. Department of Endocrinology, the First Affiliated Hospital of Wenzhou Medical University, Wenzhou 325000, China
3. Department of Laboratory Medicine, the First Affiliated Hospital of Wenzhou Medical University, Wenzhou 325000, China
4. Mass Spectrometry Lab, Instruments center for physical Science, University of Science and Technology of China, Hefei 230026, China
5. MAFLD Research Center, Department of Hepatology, the First Affiliated Hospital of Wenzhou Medical University, Wenzhou 325000, China
6. Department of Metabolic and Bariatric Surgery, The First Affiliated Hospital of Jinan University, Guangzhou 510630, China
7. Key Laboratory of Diagnosis and Treatment for The Development of Chronic Liver Disease in Zhejiang Province, Wenzhou 325000, China
8. Department of Oncology, The First Affiliated Hospital of Anhui Medical University, Hefei 230022, China
9. Inflammation and Immune Mediated Diseases Laboratory of Anhui Province, Anhui Medical University, Hefei 230032, China
10. Zhongda Hospital, School of Medicine, Advanced Institute for Life and Health, Southeast University, Nanjing 210096, China
11. Engineering Research Center of Bioprocess, Ministry of Education, Hefei University of Technology, Hefei, Anhui 230009, China
12. Lead Contact

[#]These authors contributed equally

**Table S1.** Demographics and baseline characteristics.

| | MASH patient (n=25) | Non-MASH MASLD patient (n=25) |
|---|---|---|
| Age, years | 40.56 (14.6) | 43.8 (12.9) |
| Sex | | |
| Male | 17 (68.0%) | 19 (76.0%) |
| Female | 8 (32.0%) | 6 (24.0%) |
| Race, Asian | 25 (100%) | 25 (100%) |
| Body-mass index (BMI), $kg/m^2$ | 28.7 (4.9) | 28.9 (4.2) |
| Overweight (24<BMI<28) | 5 (20%) | 8 (32%) |
| Obesity (BMI>28) | 15 (60%) | 15 (60%) |
| Body weight, kg | 81.96 (20.7) | 79.63 (15.1) |
| Body height, cm | 168.04 (9.3) | 165.44 (9.2) |
| Waist circumference, cm | 95.56 (10.5) | 93.79 (9.9) |
| Systolic blood pressure, mmHg | 134.00 (15.8) | 129.40 (11.7) |
| Diastolic blood pressure, mmHg | 87.80 (10.1) | 83.64 (6.8) |
| Fasting insulin, pmol/L | 408.52 (646.3) | 116.90 (161.7) |
| Fasting C-peptide, pmol/L | 1597.92 (1217.7) | 884.52 (870.6) |
| White blood cell, $X10^9/L$ | 6.82 (2.0) | 6.68 (1.8) |
| Red Blood Cell, $X10^{12}/L$ | 4.99 (0.5) | 4.95 (0.4) |
| Hemoglobin, g/L | 151.68 (17.6) | 150.56 (18.7) |
| Platelet, $X10^9/L$ | 260.60 (58.9) | 240.76 (60.1) |
| Total bilirubin, μmol/L | 13.68 (4.1) | 16.08 (6.3) |
| Direct bilirubin, μmol/L | 2.60 (1.2) | 2.96 (1.5) |
| Total protein, g/L | 76.06 (6.6) | 70.37 (3.5) |
| Albumin, g/L | 44.98 (4.3) | 42.08 (2.4) |
| Globulin, g/L | 31.08 (4.4) | 28.28 (2.5) |
| Albumin/Globulin (A/G) | 1.48 (0.2) | 1.50 (0.2) |
| Serum uric acid, μmol//L | 443.28 (123.3) | 409.76 (109.1) |
| Total cholesterol (TC), mmol/L | 5.24 (1.5) | 4.58 (1.0) |
| HDL, mmol/L | 1.12 (0.3) | 0.99 (0.2) |
| LDL, mmol/L | 3.38 (1.1) | 2.99 (0.7) |
| Triglycerides (TG), mmol/L | 2.11 (0.9) | 1.80 (1.0) |
| ALT, U/L | 101.72 (56.9) | 31.76 (21.8) |
| AST, U/L | 60.48 (29.1) | 29.40 (19.4) |
| ALT/AST | 1.69 (0.5) | 1.12 (0.5) |
| ALP, U/L | 83.08 (20.3) | 80.64 (24.0) |
| IgE, ng/mL | 902.22 (555.0) | 1044.19 (763.4) |

| | | |
|---|---|---|
| 5-HT, ng/mL | 1.68 (0.4) | 1.53 (0.4) |
| NEFA, mmol/L | 0.65 (0.3) | 0.57 (0.3) |

HDL high-density lipoprotein, LDL low-density lipoprotein, ALT alanine aminotransferase, AST aspartate aminotransferase, ALP alkaline phosphatase, IgE immunoglobulin E, 5-HT 5-hydroxytryptamine, NEFA Non-Esterified Fatty Acid. Data are n (%), mean (SD) or median (interquartile range).

**Table S2.** Primers for quantitative q-PCR.

| Gene | Sequence of forward primers（5' to 3'） | Sequence of reverse primers（5' to 3'） |
|---|---|---|
| *Hdc* | CGTGAATACTACCGAGCTAGAGG | ACTCGTTCAATGTCCCCAAAG |
| *Mcpt4* | TAGACCACATTCTCGCCCTTA | GGATTCTGTCTTGCTCACATCA |
| *Cma1* | TGGAGGCACGGAGTGCATA | AGGAGGACTGTTATAGACCTTCC |
| *Mcpt6* | GCCCAGCCAATCAGCG | CCAGGGCCACTTACTCTCAGA |
| *Ext1* | TGGAGGCGTGCAGTTTAGG | GAAGCGGGGCCAGAAATGA |
| *Ext2* | TGGGATCGAGGAACAAATCACC | TGCCGGTAAGTCCAGGTAGAA |
| *Ndst1* | CCACAACTATCACAAAGGCATCG | GAAAGGTGTACTTTAGGGCCAC |
| *Ndst2* | GTGGCTGATGTTGAGGCTTTG | ATCCTCCTCTTCTGTCCCGG |
| *Hpgds* | GGAAGAGCCGAAATTATTCGCT | ACCACTGCATCAGCTTGACAT |
| *Tph1* | TGACGCTGCCGATTCTCCAG | GCATGTTGCAACTCGCCAGC |
| *Cd36* | ATGGGCTGTGATCGGAACTG | GTCTTCCCAATAAGCATGTCTCC |
| *Fas* | GCTGGCATTCGTGATGGAGTCGT | AGGCCACCAGTGATGATGTAACTCT |
| *Me1* | GTCGTGCATCTCTCACAGAAG | TGAGGGCAGTTGGTTTTATCTTT |
| *Pparα* | AGAGCCCCATCTGTCCTCTC | ACTGGTAGTCTGCAAAACCAAA |
| *Pparγ* | GCATGGTGCCTTCGCTGA | TGGCATCTCTGTGTCAACCATG |
| *Atgl* | TGACCATCTGCCTTCCAGA | TGTAGGTGGCGCAAGACA |
| *Hsl* | GCGCTGGAGGAGTGTTTTT | CCGCTCTCCAGTTGAACC |
| *Mgl* | CAAGAGTGGAGCGAGCAAT | AGGACGTGATAGGCACCTTCATA |
| *Sirt1* | TGCAGACGTGGTAATGTCCAAAC | ACATCTTGGCAGTATTTGTGGTGAA |
| *Foxo1* | GCGTGCCCTACTTCAAGGATAA | TCCAGTTCCTTCATTCTGCACT |
| *Ampkα* | GTCAAAGCCGACCCAATGATA | CGTACACGCAAATAATAGGGGTT |
| *G0s2* | ACTGCACCCTAGGCCCAG | CCGAGCACCACACCGAA |
| *Htr1a* | TCAGCTACCAAGTGATCACCTCT | GTCCACTTGTTGAGCACCTG |
| *Htr1b* | TGCTCCTCATCGCCCTCTATG | CTAGCGGCCATGAGTTTCTTCTT |
| *Htr2a* | AGCTGCAGAATGCCACCAACTAT | GGGATTGGCATGGATATACCTAC |
| *Htr2b* | AAATAAGCCACCTCAACGCCT | TCCCGAAATGTCTTATTGAAGAG |
| *Htr5* | GATTGACTTCAGTGGGCTCG | AAAGTCAGGACTAGCACTCG |
| *Htr7* | CTCGGTGTGCTTTGTCAAGA | TTGGCCATACATTTCCCATT |
| *Ki67* | AATCCAACTCAAGTAAACGGGG | TTGGCTTGCTTCCATCCTCA |
| *E2f1* | CCTCGCAGATCGTCATCATC | AGAGCAGCACGTCAGAATCG |
| *Caspase 3* | TGGTGATGAAGGGGTCATTTATG | TTCGGCTTTCCAGTCAGACTC |
| *C/Ebpα* | CAAGAACAGCAACGAGTACCG | GTCACTGGTCAACTCCAGCAC |
| *Perilipin* | TGCTGGATGGAGACCTC | ACCGGCTCCATGCTCCA |
| *Leptin* | GAGACCCCTGTGTCGGTTC | CTGCGTGTGTGAAATGTCATTG |
| *Adiponectin* | GCAGAGATGGCACTCCTGGA | CCCTTCAGCTCCTGTCATTCC |
| *Srebp1* | ATCGCAAACAAGCTGACCTG | AGATCCAGGTTTGAGGTGGG |

| | | |
|---|---|---|
| *Dgat1* | ACCGCGAGTTCTACAGAGATTGGT | ACAGCTGCATTGCCATAGTTCCCT |
| *Dgat2* | AGTGGCAATGCTATCATCATCGT | TCTTCTGGACCCATCGGCCCCAGGA |
| *Gpat3* | GGAGGGCCTCGAGCTGAA | GAAGAGGTCCCCAGGAAAGC |
| *Mogat2* | CGGTCCTTCAGTGGGTCTTC | GTCTTGACCAAAGAGACAGGGA |
| *Cd11c* | CTGGATAGCCTTTCTTCTGCTG | GCACACTGTGTCCGAACTC |
| *F4/80* | CTTTGGCTATGGGCTTCCAGTC | GCAAGGAGGACAGAGTTTATCGTG |
| *β-Actin* | CATCCGTAAAGACCTCTATGCCAAC | ATGGAGCCACCGATCCACA |

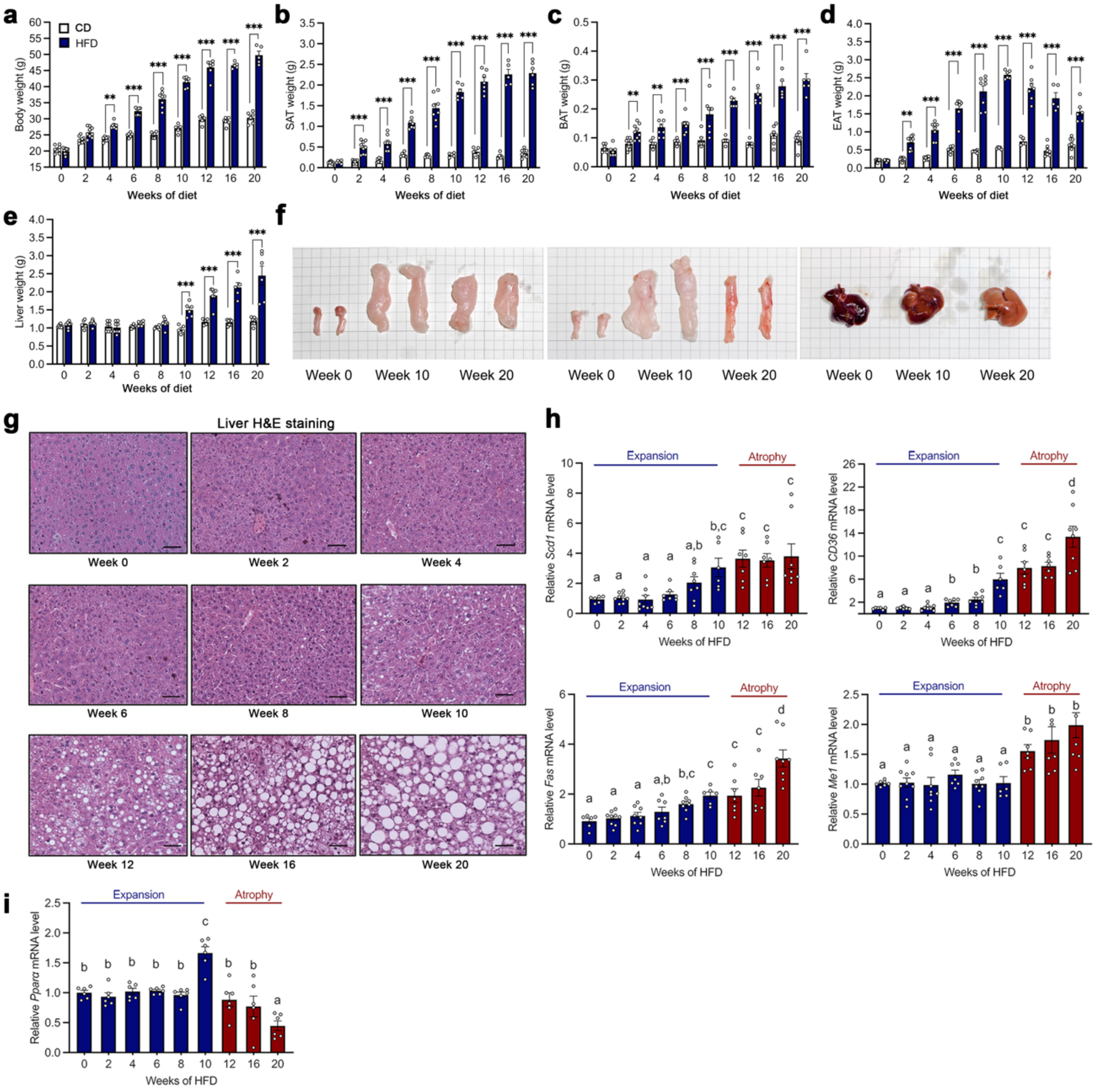


**Figure S1. HFD induces post-expansive EAT atrophy and ectopic hepatic steatosis in mice.** (**a-e**) Body weight (**a**), SAT weight (**b**), BAT weight (**c**), EAT weight (**d**), and liver weight (**e**) in WT mice on an HFD for 20 weeks. (**f**) Representative photographs of SAT, EAT and liver in WT mice fed a HFD for 0, 10 and 20 weeks. (**g**) Representative images of H&E staining in liver from WT mice on an HFD for 20 weeks. Scale bars: 50 μm. (**h**/**i**) Real-time PCR analysis of lipogenic genes stearoyl CoA desaturase 1 (*Scd1*), *CD36*, fatty acid synthase (*Fas*), malic enzyme 1 (*Me1*) (**h**), and peroxisome proliferator-activated receptor α (*Pparα*) (**i**) expression in liver from HFD-fed WT mice. Data are mean ± SEM, n=6-8/ea. Mann-Whitney U test for **a**-**e**. Kruskal-Wallis H test with Dunn's post hoc adjustment for **h** and **i**. *$p < 0.05$, **$p < 0.01$, ***$p < 0.001$. Different letters indicate statistically significant difference.

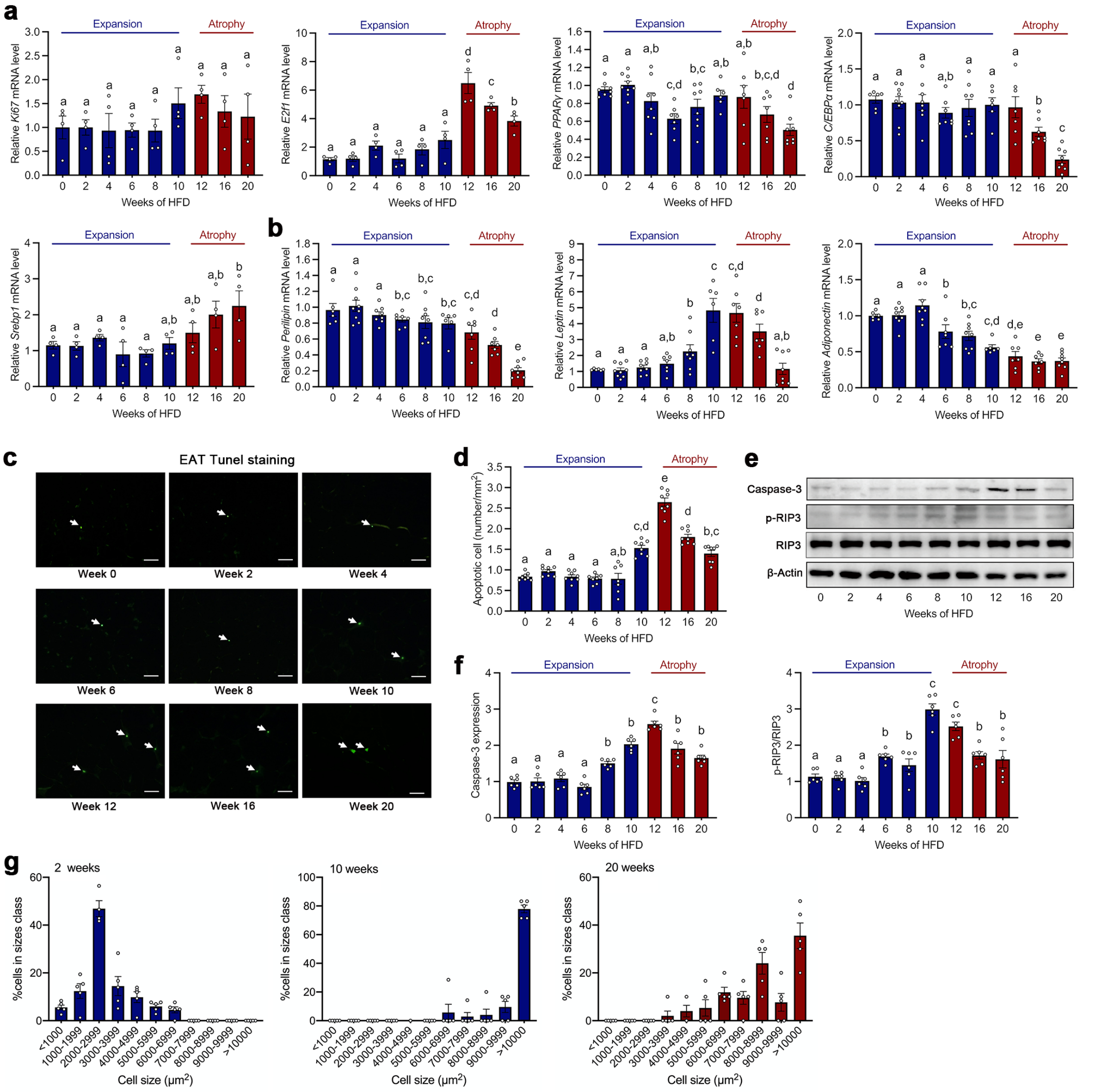


**Figure S2. Proliferation, adipogenesis, adipocyte maturation, cell death, and adipocyte size distribution in EAT during HFD feeding.** (**a/b**) Real-time PCR analysis of cell proliferation genes *Ki67* and E2F transcription factor 1 (*E2f1*), adipogenic transcription factors peroxisome proliferator-activated receptor γ (*PPARγ*), CCAAT enhancer-binding protein-α (*C/EBPα*) and Sterol regulatory element binding protein 1 (*Srebp1*) (**a**), and mature adipocyte markers *Perilipin*, *Adiponectin* and *Leptin* (**b**) expression in EAT in HFD-fed WT mice. (**c/d**) Representative images of TUNEL staining (arrows) (**c**) and TUNEL positive apoptotic cell number (**d**) in EAT in HFD-fed WT mice. Scale bar: 50 μm. (**e/f**) Immunoblot analysis (**e**) and quantification of caspase-3 to β-actin and p-RIP3 to RIP3 (**f**) in EAT in HFD-fed WT mice. (**g**) Adipocyte size and distribution in EAT in WT mice fed an HFD for 0, 10 and 20 weeks. Data are mean ± SEM, n=6-8/ea.

Kruskal-Wallis H test with Dunn's post hoc adjustment, $p < 0.05$ was considered significant. Different letters indicate statistically significant difference.

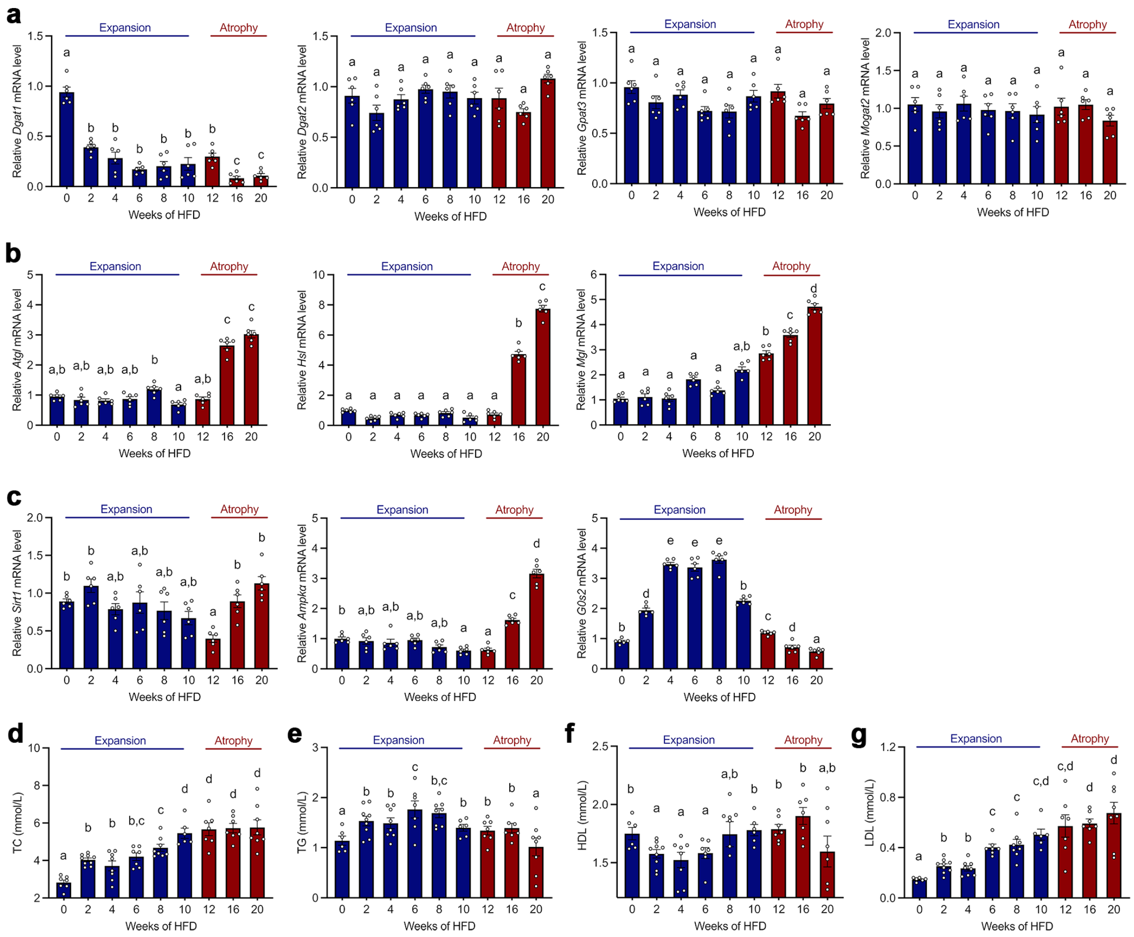


**Figure S3. Triglyceride synthesis, lipolysis, and serum lipid profile in EAT during HFD feeding.** (**a-c**) Real-time PCR analysis of triglyceride synthetic enzymes diacylglycerol acyltransferase (*Dgat*) 1, *Dgat2*, glycerol-3-phosphate acyltransferase (*Gpat3*) and monoacylglycerol acyltransferase (*Mgat2*) (**a**), lipolytic enzymes *Atgl*, hormone-sensitive lipase (*Hsl*) and monoacylglycerol lipase (*Mgl*) (**b**), and AMPK/SIRT1 signaling molecules sirtuin 1 (*Sirt1*), AMP-activated protein kinase (*Ampkα*), forkhead box O1 (*FoxO1*) and G0/G1 switch gene-2 (*G0s2*) (**c**) expression in EAT in HFD-fed WT mice. (**d-g**) Serum TC (**d**), TG (**e**), HDL (**f**) and LDL (**g**) levels in WT mice fed an HFD for 20 weeks. Data are mean ± SEM, n=6-8/ea. Kruskal-Wallis H test with Dunn's post hoc adjustment, $p < 0.05$ was considered significant. Different letters indicate statistically significant difference.

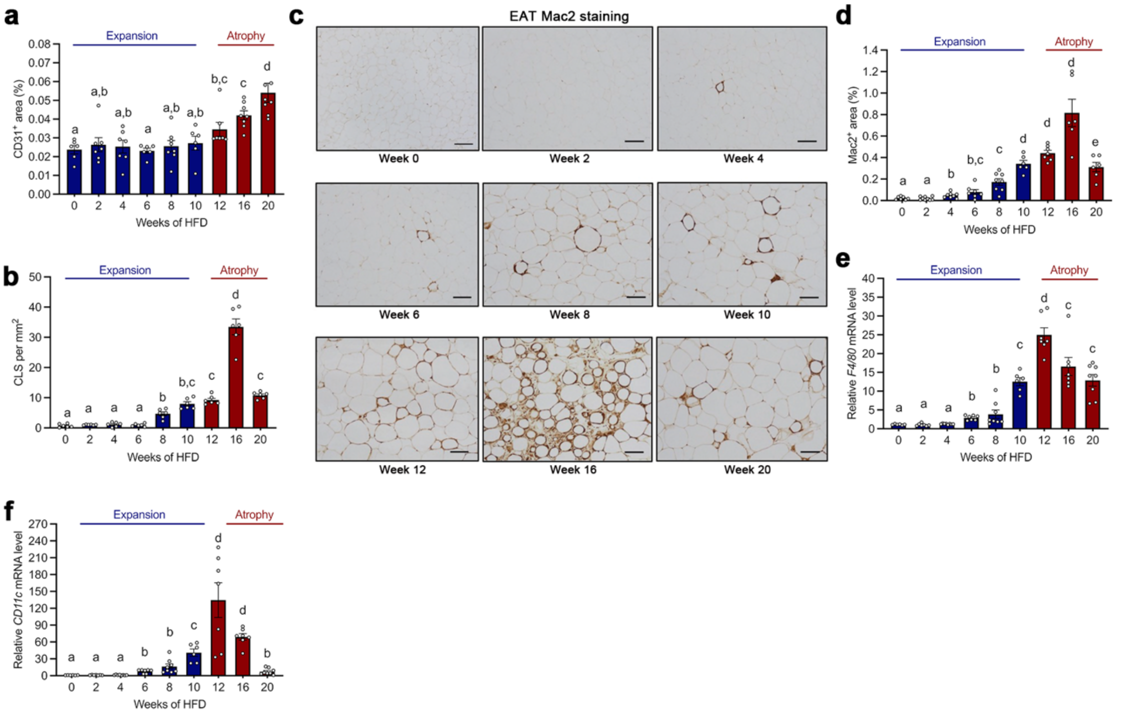


**Figure S4. HFD induces an initial increase followed by a decrease in the infiltration of pro-inflammatory macrophage during EAT atrophy.** (**a**/**b**) Quantification of CD31-positive areas (**a**) and CLS number (**b**) in WT mice fed an HFD for 20 weeks. (**c**/**d**) Representative images of Mac2 immunostaining (**c**) and Mac2-positive area (**d**) in EAT in HFD-fed WT mice. Scale bar: 50 µm. (**e**/**f**) Real-time PCR analysis of *F4/80* (**e**) and *CD11c* (**f**) expression in EAT in HFD-fed WT mice. Data are mean ± SEM, n=6-8/ea. Kruskal-Wallis H test with Dunn's post hoc adjustment, $p < 0.05$ was considered significant. Different letters indicate statistically significant difference.

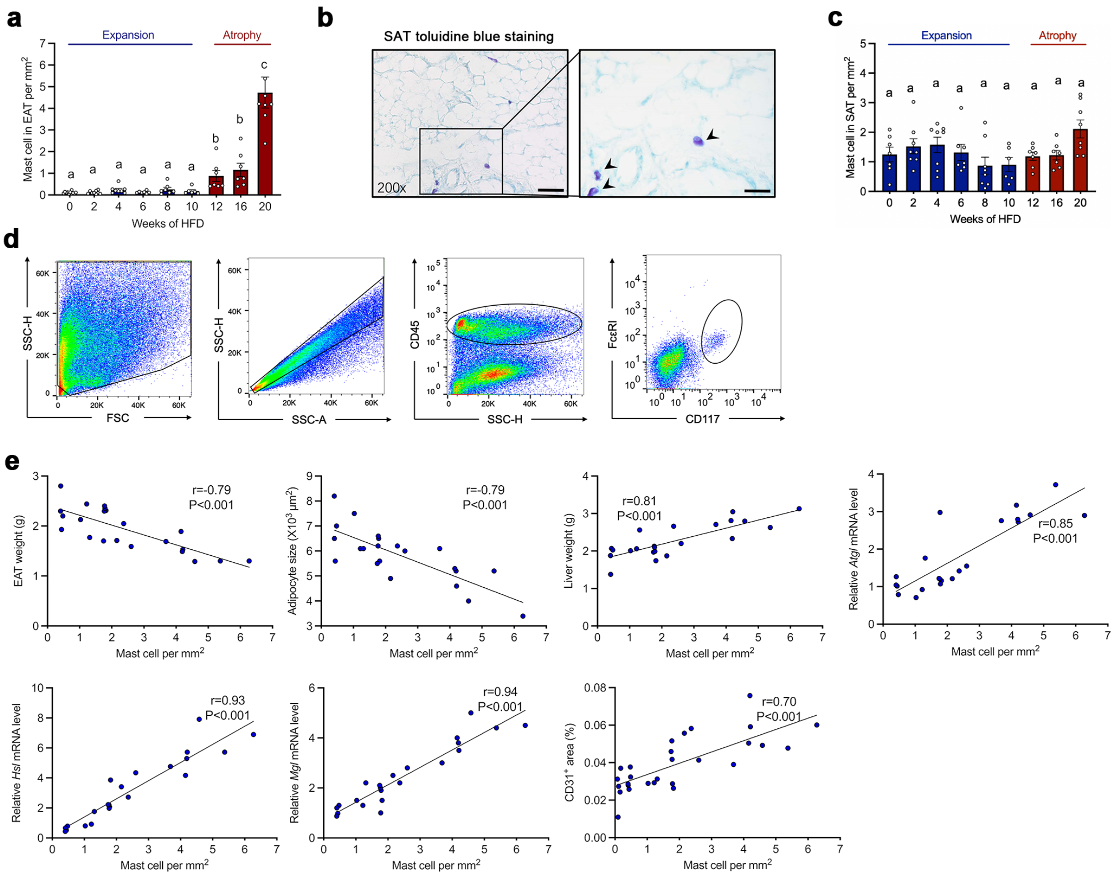


**Figure S5. MCs are accumulated during EAT atrophy in HFD-fed mice.** (**a**) MC number quantification after toluidine blue staining in WT mice fed an HFD for 20 weeks. (**b**/**c**) Representative images of toluidine blue staining for MCs (arrowheads) in SAT in WT mice at week 4 of HFD feeding (**b**) and MCs number quantification (**c**) in SAT in WT mice fed an HFD for 20 weeks. Scale bar: 50 μm, inset: 25 μm. (**d**) Gating strategy to detect MCs ($CD45^+CD117^+Fc\varepsilon RI^+$) in EAT. (**e**) Correlation tests between MC number and EAT weight, adipocyte size, liver weight, *Atgl* mRNA level, *Hsl* mRNA level, *Mgl* mRNA level or $CD31^+$ area during EAT atrophy in HFD-fed mice. Pearson's correlation test. P < 0.05 was considered significant correlation. Data are mean ± SEM, n=6-8/ea. Kruskal-Wallis H test with Dunn's post hoc adjustment for multiple testing, $p < 0.05$ was considered significant. Different letters indicate statistically significant difference.

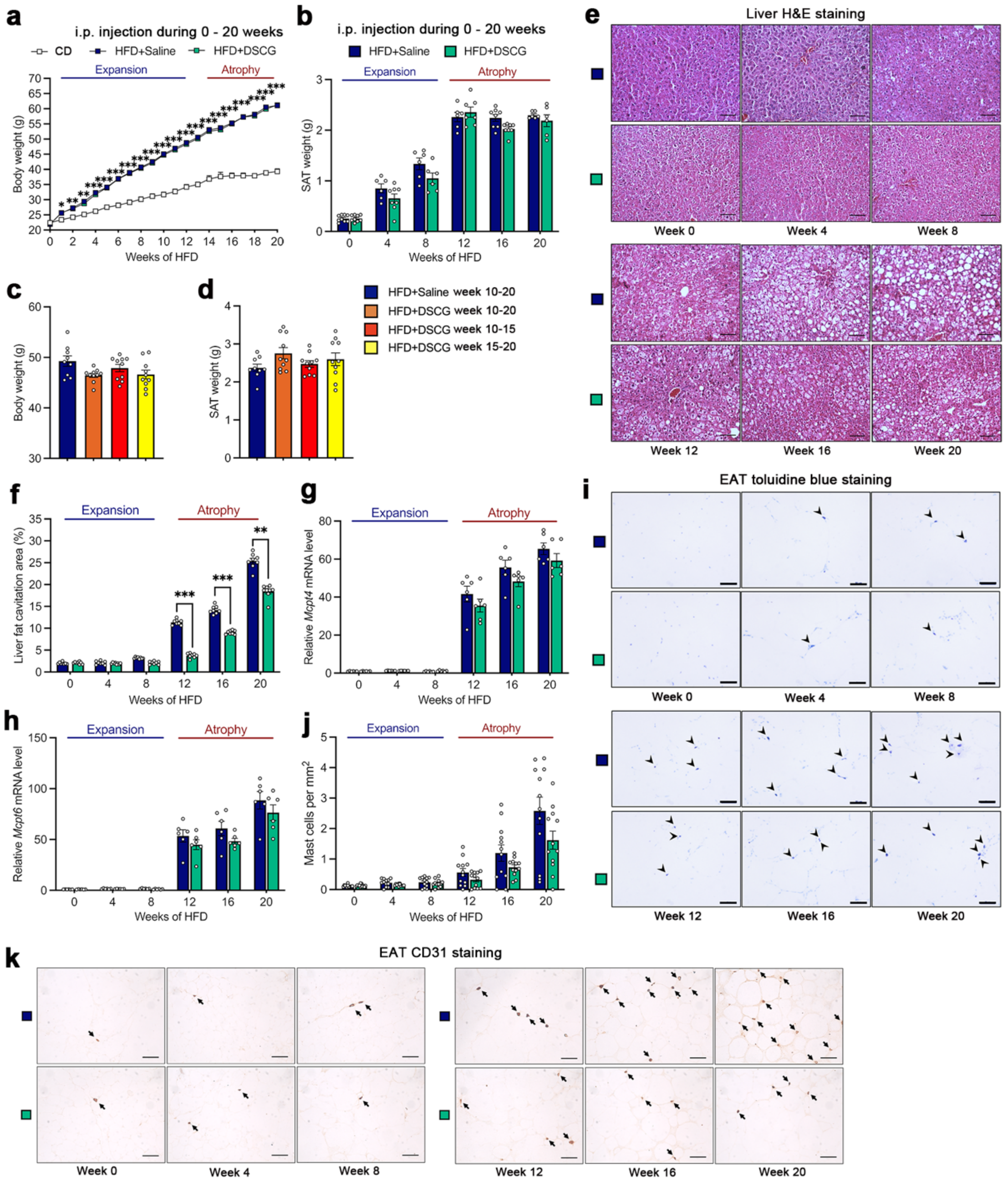


**Figure S6. MC inhibition with DSCG does not affect body weight, SAT weight and MC number, but decreases hepatic steatosis and angiogenesis in HFD-fed WT mice.** (**a**/**b**) Body weight gain (**a**) SAT weight (**b**) in WT mice fed an CD for 20 weeks, and received intraperitoneal injections of saline (blue) or DSCG (green) from week 0 to week 20 of HFD feeding. (**c**/**d**) Body weight (**c**) and SAT weight (**d**) in WT mice receiving an injection of saline (blue) or DSCG (orange) from week 10 to week 20, an injection of DSCG from week 10 to week 15 (red) or from week 15 to week 20 (yellow) of HFD feeding. (**e**/**f**) Representative images of H&E staining in liver (**e**) and percentage of liver fat cavitation area (**f**) in WT mice receiving intraperitoneal injections of saline (blue) or DSCG (green) from week 0 to week 20 of HFD feeding. Scale bars: 50 μm. (**g**/**h**) Real-time PCR analysis of *Mcpt4* (**g**) and *Mcpt6* (**h**) expression in EAT in WT mice receiving an injection of saline (blue) or DSCG (green) from week 0 to week 20 of HFD feeding. (**i**) Representative images of toluidine blue staining for MCs (arrowheads) and (**j**) MC number quantification. (**k**) Representative images of CD31 immunostaining (arrows) in EAT in WT mice receiving intraperitoneal injections of saline (blue) or DSCG (green) from week 0 to week 20 of HFD feeding. Scale bar: 50 μm. Data are mean ± SEM, n=6-8/ea. Mann-Whitney U test. *$p < 0.05$, **$p < 0.01$, ***$p < 0.001$.

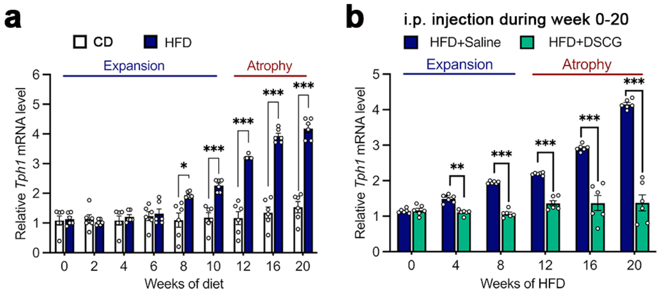


**Figure S7. *Tph1* expression in CD- or HFD-fed mice and DSCG-treated mice.** (**a**) Real-time PCR analysis of EAT *Tph1* expression in WT mice fed a CD or an HFD for 20 weeks. (**b**) Real-time PCR analysis of EAT *Tph1* expression in WT mice receiving intraperitoneal injections of saline (blue) or DSCG (green) from week 0 to week 20 of HFD feeding. Data are mean ± SEM, n=6-8/ea. Mann-Whitney U test. *$p < 0.05$, **$p < 0.01$, ***$p < 0.001$.

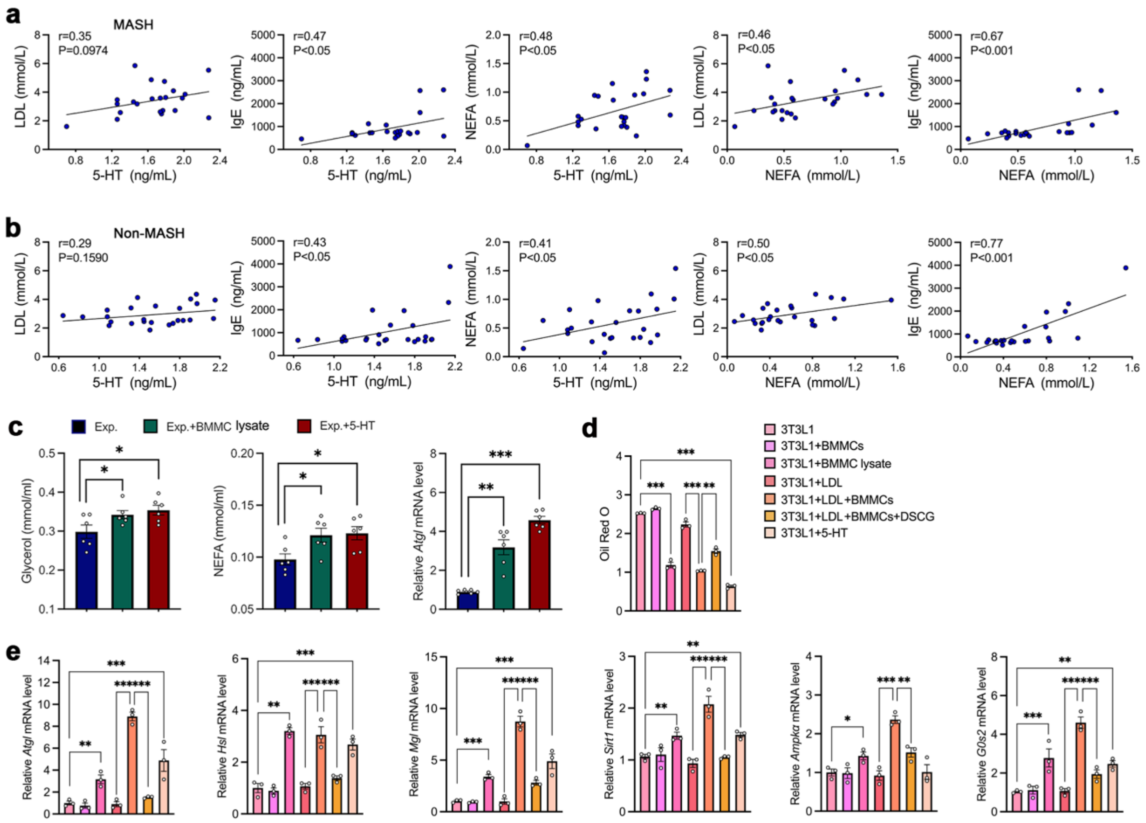


**Figure S8. MC-derived 5-HT is correlated with VAT lipolysis in MASLD patients, and promotes adipocyte lipolysis in EAT explants and 3T3L1 adipocytes.** (**a**/**b**) Serum correlation tests between LDL and 5-HT levels, IgE and 5-HT levels, 5-HT and NEFA levels, LDL and NEFA levels, and IgE and NEFA levels in MASH (**a**) and Non-MASH MASLD patients (**b**). n=50. Pearson's correlation test. P < 0.05 was considered significant correlation. (**c**) EAT explants (Exp.) from *Kit*$^{w\text{-}sh/w\text{-}sh}$ mice were cultured with or without BMMC lysate or 5-HT. Media glycerol and NEFA levels and *Atgl* mRNA level in Exp. n=5-6. (**d**/**e**) Differentiated 3T3-L1 adipocytes treated with or without live BMMCs, BMMC lysates, LDL, LDL-activated BMMCs, DSCG- and LDL-treated BMMCs, or 5-HT. Quantification of oil-red O staining (**d**), and real-time PCR analysis of *Atgl*, *Hsl*, *Mgl*, *Sirt1*, *Ampkα* and *G0s2* mRNA levels (**e**) in 3T3-L1 adipocytes. n=3/ea. Data are mean ± SEM. Welch's t-test for **c**-**e**. *$p$ < 0.05, **$p$ < 0.01, ***$p$ < 0.001.

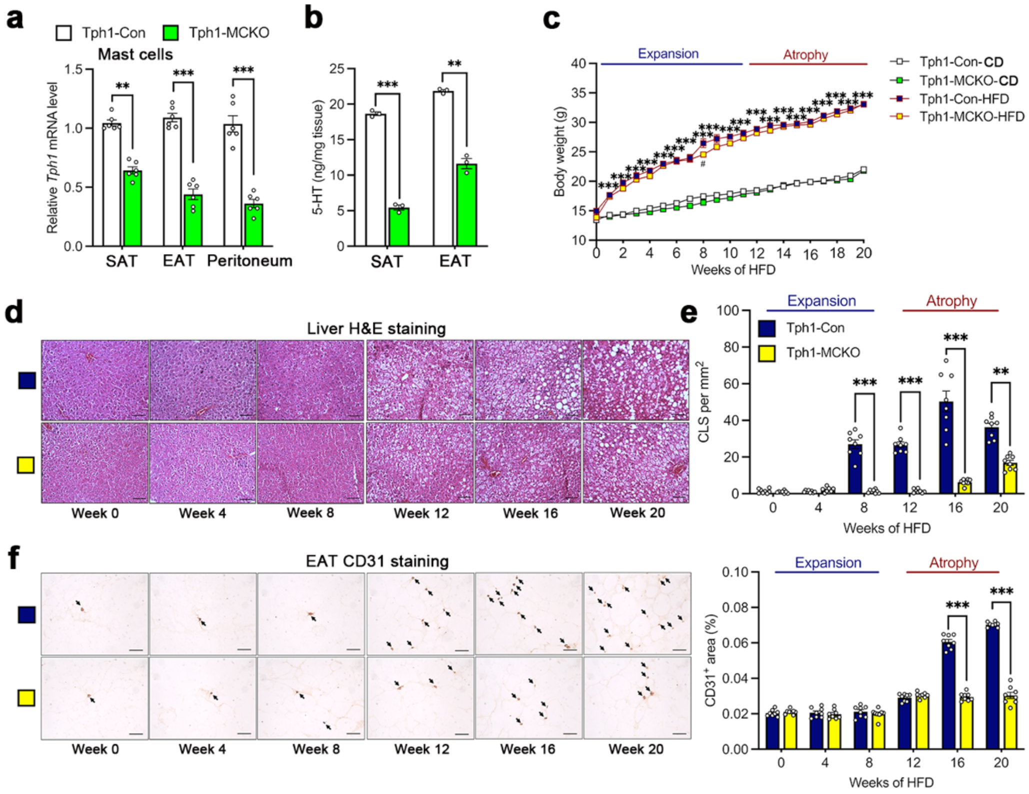


**Figure S9. MC-specific depletion of *Tph1* reduces Tph1 expression and 5-HT levels and improves hepatic steatosis and angiogenesis in HFD-fed mice.** (**a**) Real-time PCR analysis of *Tph1* expression in SAT, EAT and peritoneal MCs in CD-fed Tph1-Con and Tph1-MCKO mice. n=6/ea. (**b**) 5-HT levels in SAT and EAT in CD-fed Tph1-Con and Tph1-MCKO mice. n=3/ea. (**c**) Body weight gain in Tph1-Con and Tph1-MCKO mice during CD or HFD feeding for 20 weeks. n=6-8/ea. (**d**) Representative images of H&E staining in liver from Tph1-Con and Tph1-MCKO mice during HFD feeding for 20 weeks. Scale bar: 50 µm. n=6-8/ea. (**e**) Quantification of CLS number in EAT from Tph1-Con and Tph1-MCKO mice during HFD feeding for 20 weeks. n=6-8/ea. (**f**) Representative images of CD31 immunostaining (arrows) and CD31-positive areas in EAT in Tph1-Con and Tph1-MCKO mice during HFD feeding for 20 weeks. Scale bar: 50 µm. n=6-8/ea. Data are mean ± SEM. Mann-Whitney U test for **a**, **b**, **e** and **f**. $*p < 0.05$, $**p < 0.01$, $***p < 0.001$.

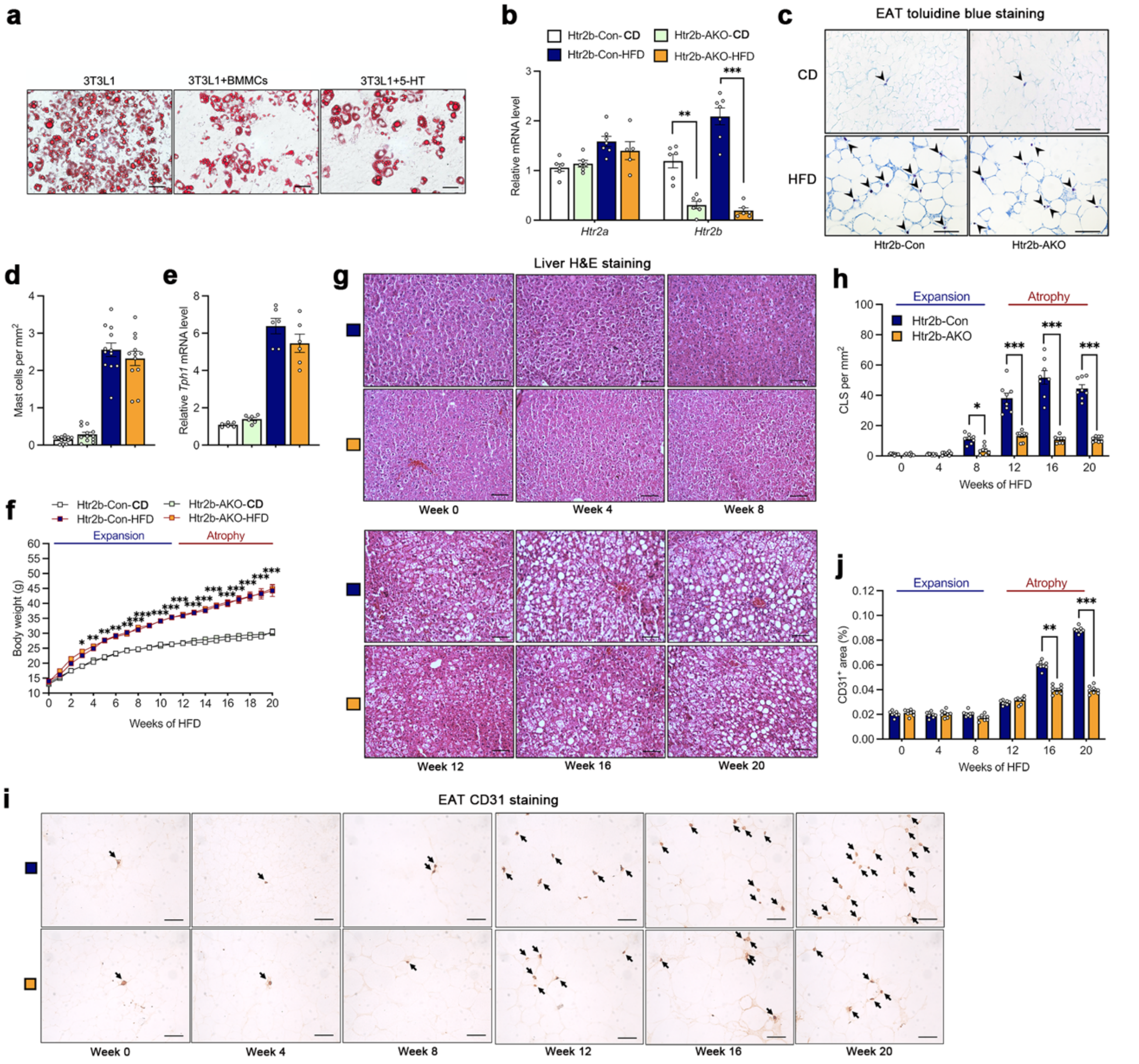


**Figure S10. Adipocyte-selective ablation of *Htr2b* does not affect MC accumulation and Tph1 expression, but reverses hepatic steatosis and angiogenesis in HFD-fed mice.** (**a**) Representative images of oil-red O staining in differentiated 3T3L1 adipocytes treated with or without BMMC lysates and 5-HT. n=3/ea. (**b**) Real-time PCR analysis of *Htr2a* and *Htr2b* expression in EAT in Htr2b-Con and Htr2b-AKO mice after HFD feeding for 20 weeks. n=6-8/ea. (**c**/**d**) Representative images of toluidine blue staining for MCs (arrowheads) (**c**) and MC number quantification (**d**) in EAT in Htr2b-Con and Htr2b-AKO mice after HFD feeding for 20 weeks. Scale bar: 50 μm. n=6-8/ea. (**e**) Real-time PCR analysis of *Tph1* expression in EAT in Htr2b-Con and Htr2b-AKO mice after HFD feeding for 20 weeks. n=6-8/ea. (**f**) Body weight gain in Htr2b-Con and Htr2b-AKO mice during CD or HFD feeding for 20 weeks. n=6-8/ea. (**g**) Representative images of H&E staining in liver in Htr2b-Con and Htr2b-AKO mice during HFD feeding for 20 weeks. Scale bars: 50 μm. n=6-8/ea. (**h**) Quantification of CLS number in EAT in Htr2b-Con and Htr2b-AKO mice during HFD feeding for 20 weeks. n=6-8/ea. (**i/j**) Representative images of CD31 immunostaining (arrows) (**i**) and CD31-positive areas (**j**) in EAT in Htr2b-Con and Htr2b-AKO mice during HFD feeding for 20 weeks. Scale bar: 50 μm. Data are mean ± SEM. Mann-Whitney U test for **b-e**, **h** and **j**.*$p < 0.05$, **$p < 0.01$, ***$p < 0.001$.
